# Deep indel mutagenesis reveals the impact of amino acid insertions and deletions on protein stability and function

**DOI:** 10.1101/2023.10.06.561180

**Authors:** Magdalena Topolska, Antoni Beltran, Ben Lehner

## Abstract

Amino acid insertions and deletions (indels) are an abundant class of genetic variants. However, compared to substitutions, the effects of indels on protein stability are not well understood and are poorly predicted. To better understand indels here we analyze new and existing large-scale deep indel mutagenesis (DIM) of structurally diverse proteins. The effects of indels on protein stability vary extensively among and within proteins and are not well predicted by existing computational methods. To address this shortcoming we present INDELi, a series of models that combine experimental or predicted substitution effects and secondary structure information to provide good prediction of the effects of indels on both protein stability and pathogenicity. Moreover, quantifying the effects of indels on protein-protein interactions suggests that insertions can be an important class of gain-of-function variants. Our results provide an overview of the impact of indels on proteins and a method to predict their effects genome-wide.

## Introduction

Short insertions and deletions (indels) are an abundant source of genetic variation^1–4^, accounting for ∼12% of all variants in the human genome^5^. Indels are more challenging to accurately genotype than substitutions^6,7^ and their effects are less well understood, even in protein coding regions ^8–14^. Indeed many of the most popular variant effect prediction (VEP) algorithms do not even return scores for indels^12^ (Extended Data Table 1) and the performance of those that do is unclear due to a lack of benchmark data^15^.

Indels are a fundamentally different type of perturbation to substitutions. Whereas substitutions are ‘side chain mutations’, indels are ‘backbone mutations’ that can have a much more severe effect on structure, stability and function^16–19^. The most frequent indels - accounting for about half of indels^6,20^ - in natural genomes are change in copy count (CCC) variants, where one or more nucleotides are repeated or deleted, most often due polymerase slippage during DNA replication^21,22^.

Massively parallel DNA synthesis-selection-sequencing experiments - also called deep mutational scanning (DMS) or multiplex assays of variant effect effects (MAVEs) - provide a great opportunity to quantify the effects of indels on proteins at scale in order to better understand their effects and to evaluate and develop computational methods for indel variant effect prediction^23,24^. Pioneering studies on individual proteins^8–14^ have recently been supplemented by the generation of a much larger dataset where the effect of indels on *in vitro* protein fold stability were quantified using a protease-sensitivity assay^25^. Here we use this dataset, as well as new deep indel mutagenesis quantifying *in vivo* protein abundance across structurally diverse domains to test the generality of previously proposed and new hypotheses for indel tolerance. We also use the data to develop a series of models – INDELi - that perform well for predicting the effects of indels on protein stability. These models also perform well for predicting indels that are pathogenic in humans, consistent with reduced protein stability being the major mechanism by which indel variants cause genetic diseases.

## Results

### Deep Indel Mutagenesis of *in vivo* protein domain abundance

To better understand and predict the impact of insertions and deletions on proteins, we first performed deep indel mutagenesis (DIM) of structurally diverse protein domains. For each domain, we designed a library of variants covering all sequential deletions of 1 to 3 aa, all change in copy count (CCC) insertions that repeat 1-3 aa, and all 3 nt out-of-frame deletions that remove one aa and substitute the next in a single event (delSubs) (Fig. 1a). We quantified the effect of each variant on the abundance of each domain using a highly-validated protein fragment complementation assay (PCA), abundancePCA (aPCA).^26–28^. Briefly, the variant library is expressed in *Saccharomyces cerevisiae* fused to a fragment of a reporter enzyme (dihydrofolate reductase; DHFR) such that the cellular growth rate quantifies abundance over at least three orders of magnitude^26,27^. All transformation-selection-sequencing experiments were performed in triplicate and were highly reproducible (Extended Data Fig. 1a, Pearson’s r=0.920-0.928). aPCA substitution and indel scores also correlated well with *in vitro* biophysical measurements of fold stability^25,29^ (Fig. 1b, r=0.564-0.920 and Extended Data Fig. 1b, r=0.410-0.858).

**Figure 1.**
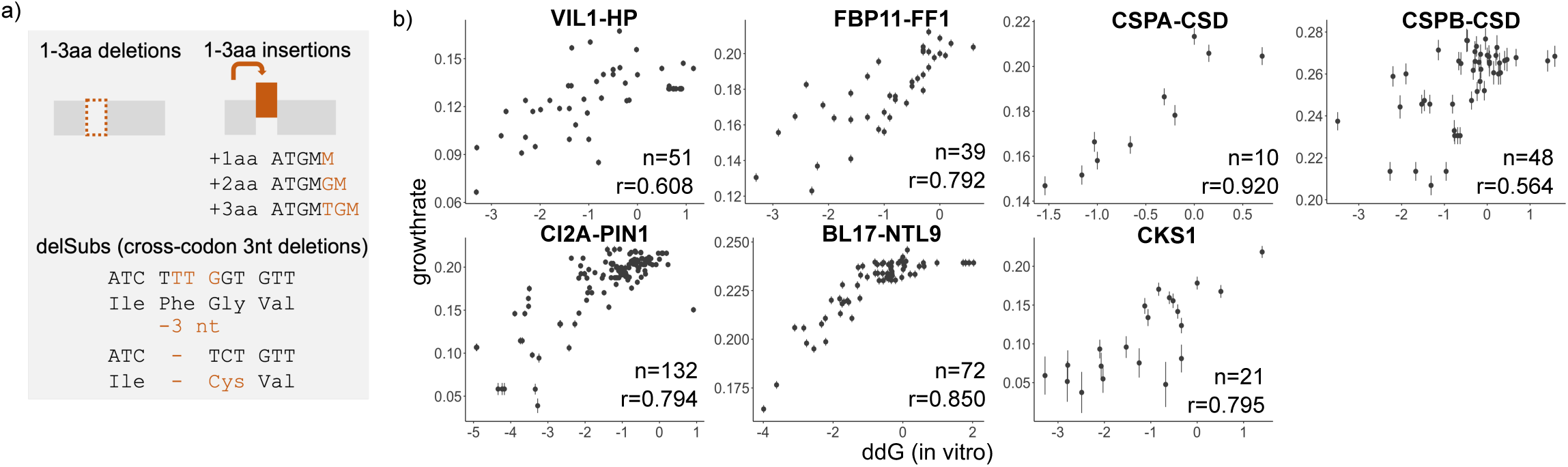
Deep indel mutagenesis of diverse protein domains. **a.** Library design for the 9 protein domains, including single and multi-aa indels, delSubs and single aa substitutions. **b.** Correlation between *in vitro* measured ddG stability and the aPCA scores of selected substitution mutants. Selection of mutants was based on available *in vitro* ddG scores in Protherm^29^.

### Patterns of indel tolerance within and among domains

All nine protein domains are highly intolerant of aa deletions, with 70% of 1aa deletions strongly reducing abundance (abundance < -0.546, variants within one standard deviation of the deleterious mode, Extended Data Fig. 1c). In general, 1aa CCC insertions are slightly better tolerated (Kolmogorov-Smirnov test, p-value=0.0343), with 64% strongly reducing abundance (Extended Data Fig. 1c). Overall, indel tolerance varies across domains. For instance, domains like FBP11-FF1 and GRB2-SH3, are almost entirely intolerant of 1aa indels, only tolerating them in domain termini (Figure 2a, 81% and 83% of 1aa indels are severely deleterious). In contrast, CSPB-CSD and PSD95-PDZ3, can accommodate 1aa indels throughout their tertiary structure, with only 54% and 53% of indels being severely deleterious.

**Figure 2.**
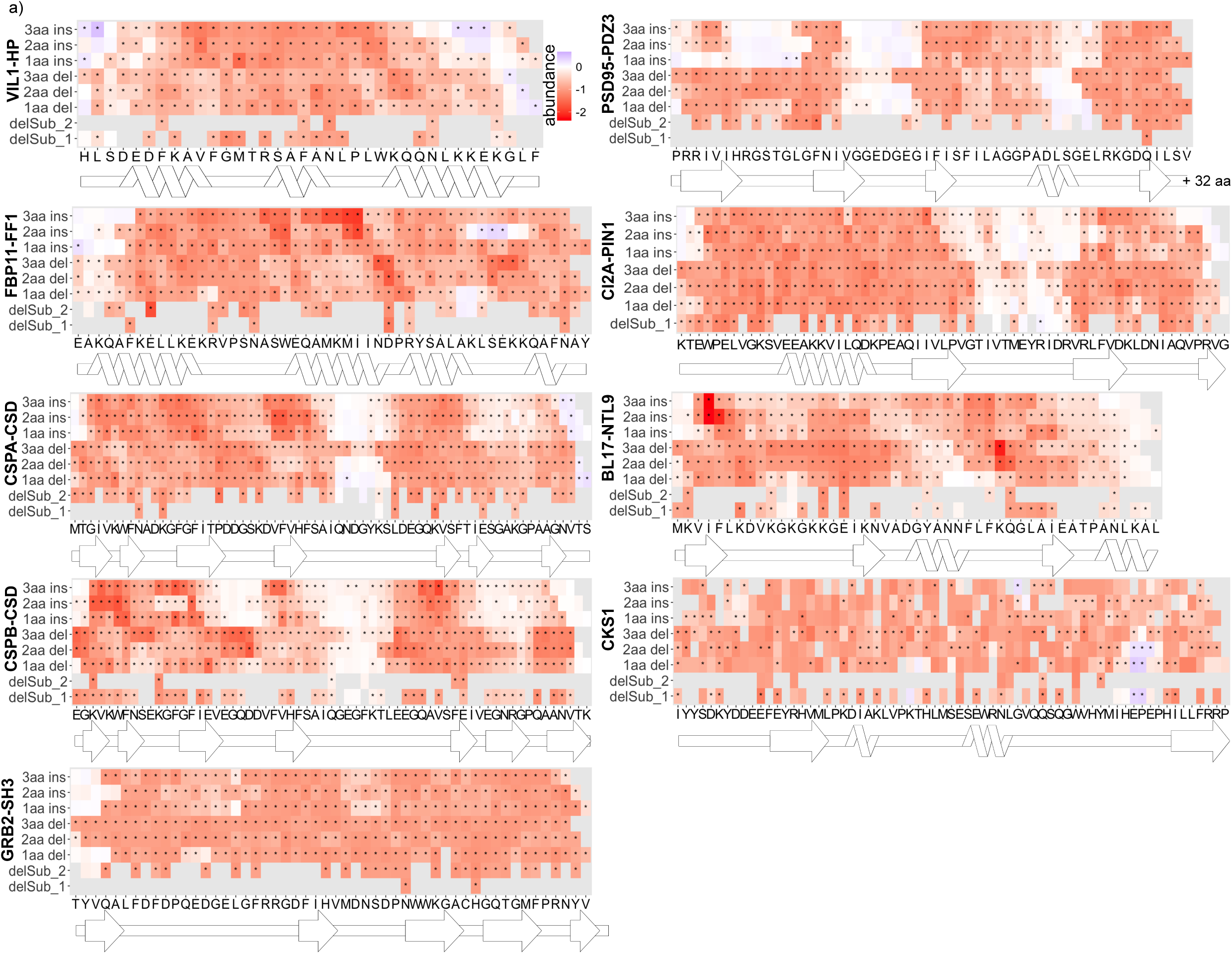
Patterns of indel tolerance within and between domains. **a.** Heat maps of protein domain abundance effects of all mutation types with significant changes in abundance (Bonferroni multiple testing correction of z-stats in a two-tailed test) marked with “*”. We show the 2-dimensional structural representation of the domains (SSDraw^63^) under each heatmap. For PSD95-PDZ3 and GRB2-SH3 only, we mutated the first 52 residues, leaving, respectively, 32 and 4aa in the C-terminal as wild type.

The relative tolerance to 1aa indels also varies within each protein, with regions similarly tolerant to insertions and deletions, regions more tolerant to 1aa deletion, and regions more tolerant to insertion (Fig. 2a). For example, in BL17-NTL9, 1aa indels are similarly tolerated in helix 2 while in helix 1 of the same protein, 1aa deletions are less detrimental than insertions. In CSPB-CSD, insertions and deletions are similarly well-tolerated in loop 4, whereas in loop 2, insertions are more detrimental than deletions, and in loops 3, 5 and 6 deletions are more detrimental than insertions (Fig. 2a). In the recent mutagenesis of the Kir2.1 potassium channel, longer (2 and 3 aa) deletions and insertions are more detrimental than shorter ones (1aa)^12^. Considering all nine proteins, 3aa deletions are indeed more detrimental than 1aa deletions, but this is not true for insertions (Wilcox one-sided t-test, Extended Data Fig. 1e and i-l). Moreover, this again varies quantitatively across proteins and in different regions of the same protein (Fig. 2a). For example, in CI2A-PIN1, 2-3 aa indels are tolerated similarly to 1aa indels across the entire domain, while in BL17-NTL9 longer indels are more detrimental in certain regions, like strand 1 or loop 3 (Fig. 2a). Considering the entire dataset, effects of 1aa CCC insertions are reasonably correlated to the effects of 1aa deletions (r=0.484, p-value<2.2e-16, Extended Data Fig. 1f), but the effects of 2aa and 3aa CCC insertions are less well correlated with the effects of 2aa and 3aa deletions (2aa: r=0.279, p-value=2.4e-9; 3aa: r=0.145, p-value=0.00224; Extended Data Fig. 1g and 1h)).

Approximately 1% of indels increase protein domain abundance above wild type levels. These are mainly located at the N- or C-termini of the protein domains (Fig. 2a) with the exceptions of FBP11-FF1 where 2aa insertions increase domain abundance in loop 3 and CKS1 where 1-2aa deletions increase abundance at the end of loop 3.

### Contrasting the impact of insertions and deletions to substitutions

For two of the proteins, PSD95-PDZ3 and GRB2-SH3, in addition to the CCC insertion repeats, we also quantified the effects of all 20 aa insertions before every position and the effects of all 19 aa substitutions at every site, allowing us to directly compare the effects of inserting and substituting to the same residues (Fig. 3a). For PSD95-PDZ3, which tolerates indels throughout its tertiary structure (Fig. 3a), the average effect of substitutions predicts reasonably well the effect of deleting the residue (r=0.534, Fig. 3b) as well as CCC insertions after (r=0.441, Fig. 3b) and before (r=0.313, Extended Data Fig. 2b) the site. For GRB2-SH3, which is highly intolerant of indels (Fig. 3a), substitution tolerance is reasonably predictive of CCC insertions after the substitution site (r=0.397) but less predictive of the tolerance for insertions before a site (r=0.0617 [p=0.667], Extended Data Fig. 2b) and deletion tolerance (r=0.208 [p=0.143], Fig. 3b).

**Figure 3.**
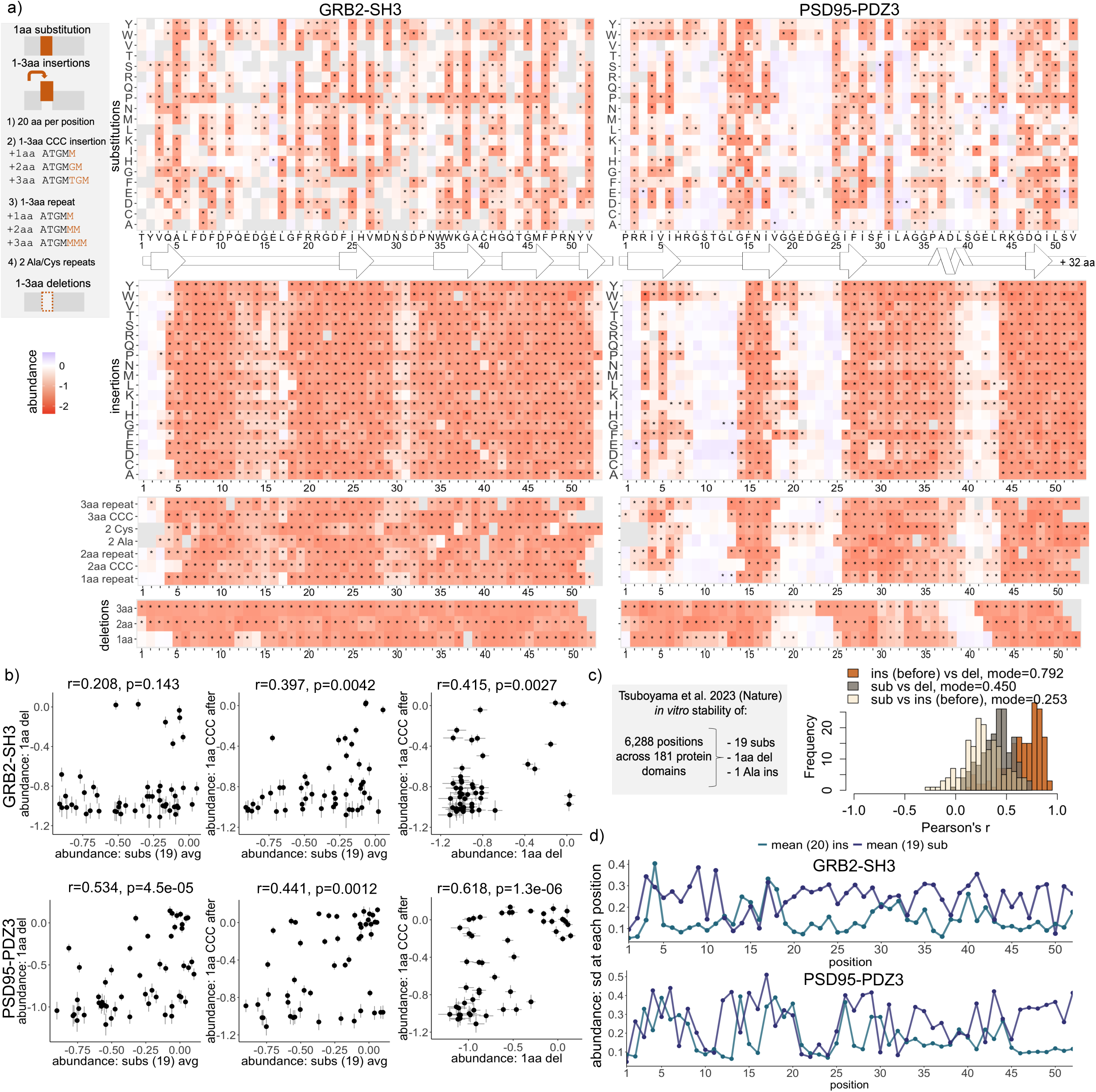
Contrasting the impact of insertions and deletions to substitutions. **a.** Heatmaps of 19 single aa substitutions, 20 aa insertions, 1-3aa insertion repeats and 1-3aa deletion effects on protein abundance. Changes in abundance significant from the weighted mean of the synonymous variants are marked with “*” (Bonferroni multiple testing correction of z-stats in a two-tailed test); y-axis: identity, type or length of mutation; x-axis: mutated position- and the aa sequence of the domain. We show the 2-dimensional structural representation of the domains (SSDraw^63^) under each heatmap. For PSD95-PDZ3 and GRB2-SH3, we mutated the first 52 residues, leaving, respectively, 32 and 4aa in the C-terminal as wild type. **b.** Scatter plots of 1aa deletion or 1aa CCC insertion versus effects of mean substitution/residue with Pearson’s correlations and corresponding p-values. Error bars represent the replicate standard deviation from the aPCA assay. **c.** Overview of the filtered Tsuboyama et al. dataset (See Methods) and histograms of correlations of indel and substitution effects for 171 domains (>9 indels) from Tsuboyama et al. The modes of Pearson correlations are displayed in the legend. **c.** Variation of 19 substitution (purple) and 20 insertion (blue) effects per position as standard deviation of effects/residue.

We next considered the large-scale mutagenesis dataset of Tsuobyama et al. ^25^, which used a protease-sensitivity assay to quantify the effects of variants on *in vitro* fold stability. Filtering the high-confidence subset of this dataset for positions with measurements of 1aa deletions, 1 Ala insertions and 19 substitutions, results in a dataset of substitutions and indels at 6,288 positions across 181 protein domains (see Methods and Fig. 3c).

Across 171 protein domains (domains with >9 indel measurements), substitution tolerance predicts Ala insertion tolerance similarly well after (mode r=0.263) and before a residue (r=0.253) and better predicts 1 aa deletion tolerance (mode r=0.450, Fig. 3c and Extended Data Fig. 2c). Interestingly, substitution tolerance is a better predictor of insertion and deletion tolerance in loops than in secondary structure elements (mode r=0.510 and r=0.544 for insertions and deletions in loops, mode r=0.219 and r=0.333 for insertions and deletions in secondary structure elements, Extended Data Fig. 2d). That substitution tolerance predicts indel tolerance reasonably well - and variably so in different protein regions - suggests a strategy for predicting indel variant effects (see below).

### Insertions of different amino acids

Although CCC insertions are by far the most frequent aa insertions in natural genomes^6^, experimentally we can insert any aa at any position. Substitutions to different aa often have strikingly different effects at the same residue^30^ (Fig. 3a and Extended Data Fig. 2e). Such variation is less frequent for 1aa insertions, with a lower standard deviation of variant effects at most positions (Fig. 3d). However, this again varies across sites, for example, at positions 4 and 15 in GRB2-SH3 and positions 14 and 20 in PSD95-PDZ3 insertions have a wide range of effects (Extended Data Fig. 2e). At other sites, most insertions are tolerated and only particular aa are detrimental. Examples include positions 17 and 53 in GRB2-SH3 and positions 39 and 42 in PSD95-PDZ3 (Extended Data Fig. 2e). At other sites most 1aa insertions are deleterious while specific aa are tolerated, for example positions 35 and 40 in GRB2-SH3 (Extended Data Fig. 2e).

Considering all sites, the average effect of inserting an aa is correlated to the average effect of substituting to the same aa (r=0.606, p-value=0.0046 and r=0.551, p=0.0119 for GRB2-SH3 and PSD95-PDZ3), although the effects of insertion are typically much more detrimental (Extended Data Fig. 2f,g).

### DelSubs

Out of frame 3nt deletions can cause a more complicated protein sequence change where one aa is deleted and the next aa is substituted. We refer to these variants as ‘delSubs’. An important example of a pathogenic delSub is the F508del variant in the cystic fibrosis transmembrane conductance regulator (CFTR) gene, which is the most frequent cause of cystic fibrosis^31^.

To better understand delSubs, we quantified the effects of 325 across the nine structurally diverse proteins. Our data shows that the effects of delSubs correlate well with the effects of 1aa deletions of the same residues (Fig 4; r=0.729, p-value<2.2e-16), showing that delSub tolerance is predominantly driven by 1aa deletion tolerance. However, for a subset of sites where 1aa deletions are tolerated, delSubs are detrimental. At these sites, which constitute ∼8% of delSubs, the additional substitution results in destabilization of the protein (Fig. 4).

**Figure 4.**
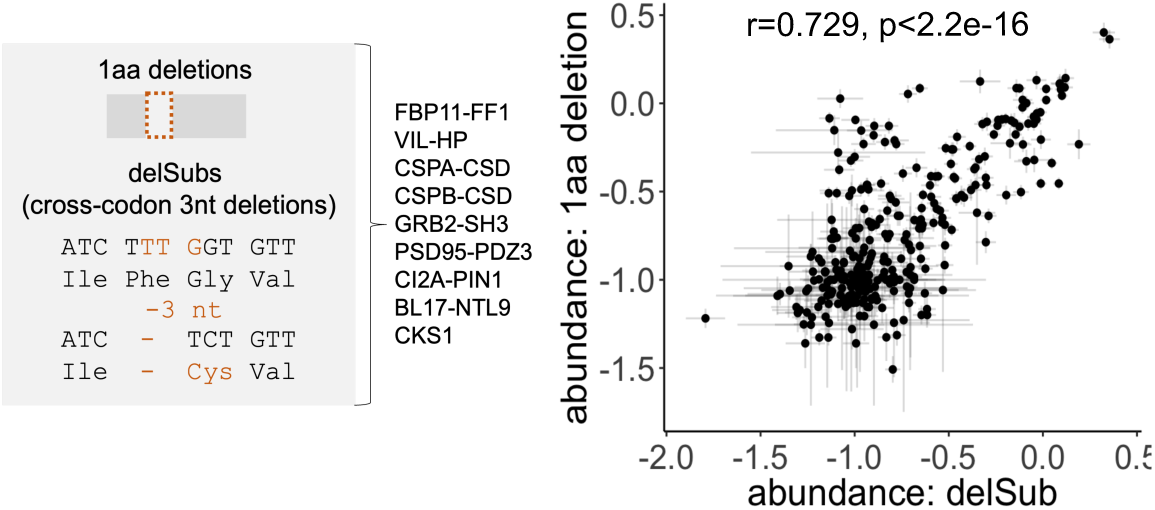
Tolerance of DelSubs. Scatter plot of 1aa deletions and 1aa delSub abundance scores for 9 protein domains and the corresponding Pearson’s correlation coefficient.

### Structural determinants of indel tolerance

Previous studies on individual proteins have proposed many different hypotheses for structural determinants of insertion and deletion tolerance^8–10,12,13^. The quantification of thousands of insertion, deletion and substitution variants across diverse protein folds by Tsuobyama et al.^25^ provides an opportunity to better understand how their effects relate to structure.

Mutations, in general, have been proposed to be more detrimental in the buried hydrophobic cores of proteins^10^. However, analyzing substitutions, 1 Ala insertions and 1aa deletions across 171 domains shows that solvent exposure is a much less important determinant of indel tolerance than of substitution tolerance. Whereas relative solvent accessibility (rSASA) alone is a good predictor of substitution tolerance (mode r=0.644, Fig. 5a and Extended Data Fig. 3a), it only moderately predicts deletion tolerance (mode r=0.364) and poorly predicts insertion tolerance (mode r=0.168). The importance of solvent accessibility in determining substitution tolerance is further illustrated by the periodic patterns of substitution tolerance in secondary structure elements. In alpha-helices, substitution tolerance follows the 3-4 aa structural periodicity of helices, consistent with substitutions of side chains on the same buried face of a helix having more similar effects (Fig. 5b). Similarly, in beta-strands, substitutions in every second aa have more similar effects, consistent with their side chains being on the same face of a strand (Fig. 5b). Neither of these periodicities in variant effects is observed for insertions or deletions (Fig. 5b and Extended Data Fig. 3b). These differences likely reflect the fundamental difference between substitutions, which are side-chain perturbations, and insertions and deletions, which are backbone perturbations. Structural analyses have suggested that proteins can locally accommodate insertions in surface residues of helices and strands by pushing the indel out as a “bulge”, preserving the structural register^1,32,33^. Our analyses suggest that this is a rare occurrence.

**Figure 5.**
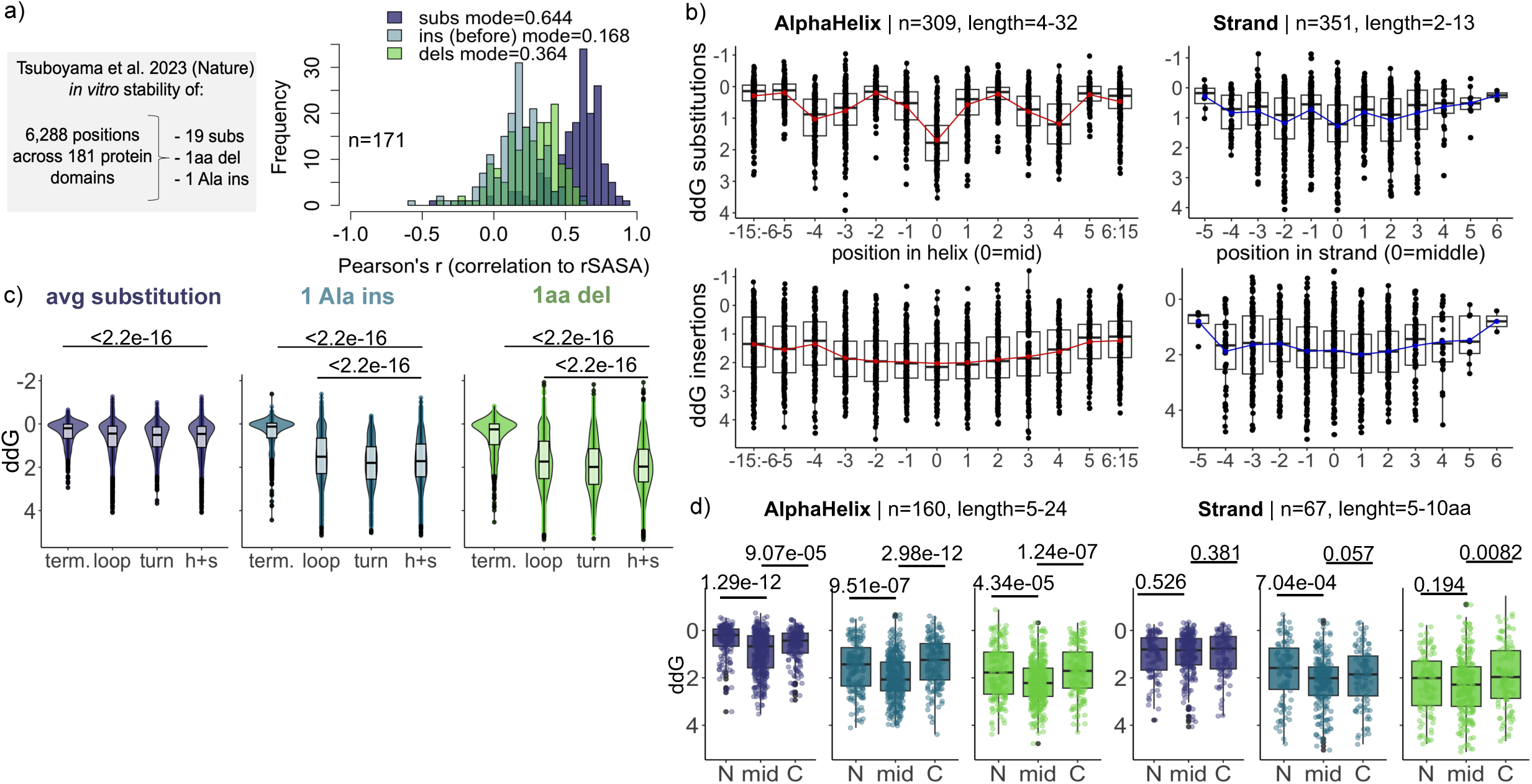
Structural determinants of indel and substitution tolerance. **a.** Overview of the filtered Tsuboyama et al. dataset (See Methods) used for analysis throughout this figure, and histograms of Pearson’s correlation coefficients of 1aa deletions, 1 Ala insertions or average substitution stability ddG and rSASA. The mode of Pearson correlations are displayed in the legend. **b.** Patterns of ddG periodicity for substitutions and 1 Ala insertions across alpha-helices (red) and strands (blue). Red/blue line represents the mean ddG/position. Position in helix/strand on the x-axis is counted from the middle of the structural element (See Methods). **c.** Violin plots of average substitution, 1 Ala insertion and 1aa deletion ddG across domain termini (term.), loops, turns between strands and helices and strands (h+s). P-values from Wilcox one-sided t-tests are displayed above the plots. **e.** Box plots of average substitution, 1 Ala insertion and 1aa deletion ddG across the N-terminal (N), middle (mid) and C-terminal (C) residues of alpha-helices and strands. N- and C-termini are defined as the most terminal residues of the helix/strand, while the midpoint is designated as position Additional data points at positions +1 or -1 relative to position 0 are included, if available.

Mutagenesis of the potassium channel Kir2.1^12^ suggested that N- and C-termini are more tolerant of substitutions and indels compared to structured regions. Across 181 domains, N- and C-termini are indeed more tolerant of substitutions, insertions and deletions than helices and strands (Figure 5c). Moreover, short termini (<= 3aa) generally display high tolerance to substitutions and indels, while long termini (≥4 aa) are less tolerant, with insertions and deletions being more destabilizing at 3-4 positions before and after secondary structure in N- and C-termini respectively (Extended Data Fig. 3e-f). Substitutions are typically weakly destabilizing in the 1-2 positions before secondary structure in long N-termini and in the position immediately after secondary structure in C-termini (Extended Data Fig. 3e-f).

It has also been previously reported in some proteins that indels are better tolerated in loops between secondary structure elements^8,9,12^. Across 181 protein domains, loops, excluding turns between strands, are indeed more tolerant of of deletions (Figure 5c, Cohen’s d effect size=-0.194, p-value<2.2e-16) and insertions (effect size=-0.184, p-value<2.2e-16) than structured regions. However, insertions and deletions are just as deleterious across turns between strands as they are in structured elements (Figure 5c). Neither of these differences are observed for substitutions (Figure 5c).

Mutagenesis of Kir2.1 also suggested that beta strands are more sensitive to indels than helices^12^. Although this conclusion is supported across 181 domains, the differences are small (effect size=-0.172, p-value=5.42e-5 for 1aa deletions and effect size=-0.117 and p-value=0.0108 for 1 Ala insertions, Extended Data Fig. 3c) and only significant for substitutions when restricting the analysis to proteins containing both helices and strands (n=45, Extended Data Fig 3d). Finally, mutagenesis of the TEM-1 β-lactamase suggested that the ends of secondary structure elements are more tolerant to indels than the middle residues^9^. Across 181 domains, both ends of alpha-helices are indeed slightly more tolerant of insertions, deletions and substitutions (Fig. 5d). However, the ends of beta-strands are not more tolerant of substitutions, and insertion and deletion tolerance is only very weakly and asymmetrically increased (Fig. 5d).

### Evaluating indel variant effect prediction

Large experimental testing of indels provides the first opportunity to systematically evaluate computational indel variant effect predictors (VEP). We evaluated two widely used VEPs that return indel predictions, CADD^34^ and PROVEAN^35^, and the large language model ESM1b^36^ which was recently adapted for indel prediction^37^

CADD only provides predictions for human proteins and its performance is reasonable for substitutions (mode r=0.357) but very poor for insertions (mode r=-0.071) and deletions (mode r=-0.082, Fig. 6a). PROVEAN’s performance is slightly worse for substitutions (mode r=0.287) but better for deletions (mode r=0.324) and insertions (mode r=0.428). The performance also varies extensively across proteins (interquartile range r=0.259-0.582 for insertions and r=0.260-0.570 for deletions). ESM1b, in contrast, provides reasonable prediction of substitution (mode r=0.502), insertion (mode r=0.547) and deletion (mode r=0.575) variant effects. However, performance again varies extensively across domains (Fig. 6a, e.g. interquartile range r=0.188-0.609 for insertions and r=0.274-0.642 for deletions).

**Figure 6.**
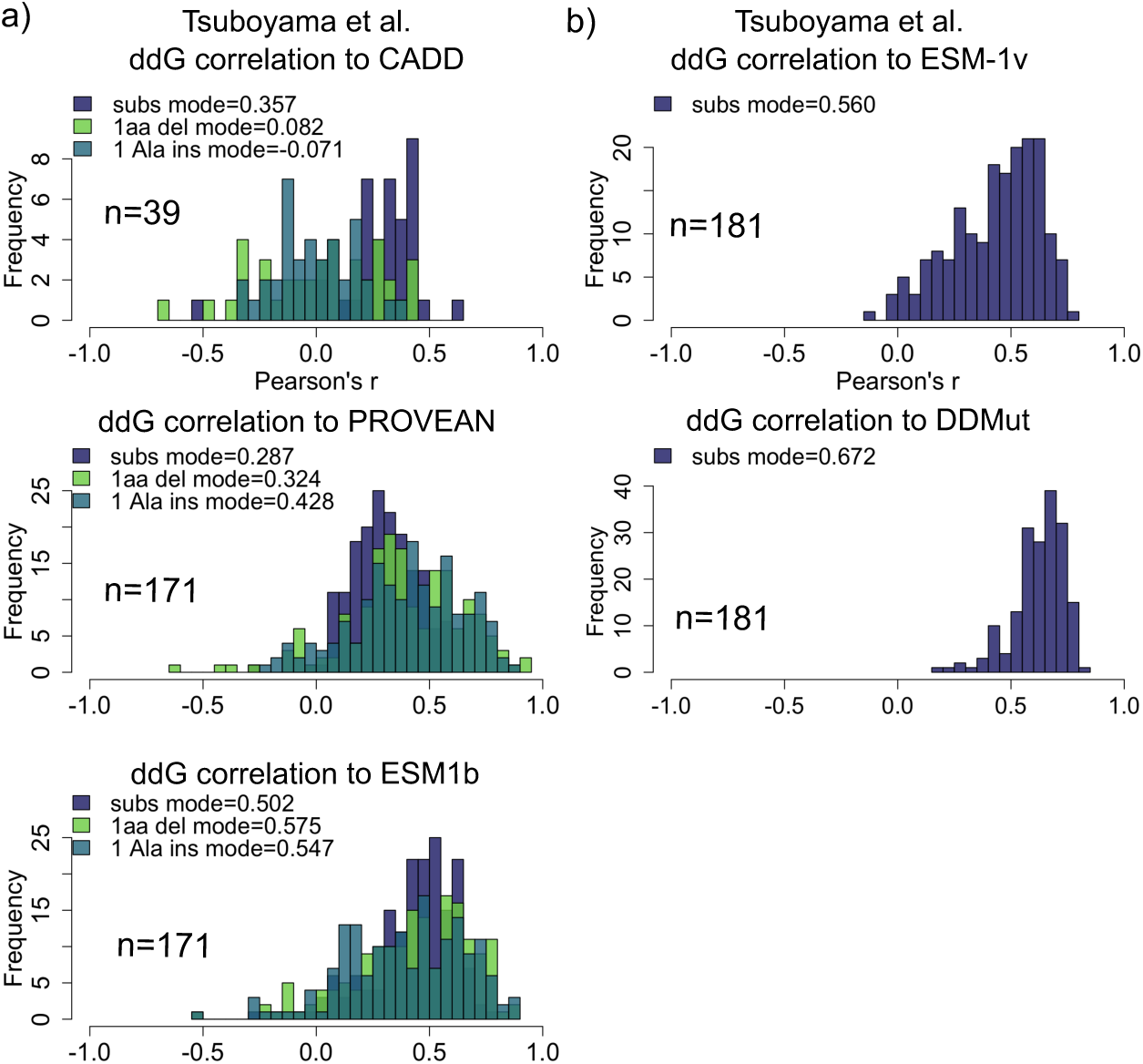
Evaluating indel variant effect prediction. **a.** Histograms of Pearson’s correlations coefficients for observed and predicted tolerance scores (>9 indel scores) of average substitutions (purple), 1 Ala insertions (blue) and 1aa deletions (green) across protein domains from Tsuboyama et al.. Prediction accuracy of CADD^34^ could only be tested on human domains. **b.** Histograms of Pearson’s correlations coefficients for observed and predicted tolerance scores of all substitutions (purple), predicted using ESM-1v^38^ and DDMut^40^.

We also evaluated the performance of state-of-the-art VEPs that do not return scores for indels for predicting the effects of substitutions on fold stability and found that they can be reasonably good. For example, mode r=0.560 for the large language model ESM-1v^38^, mode r=0.610 for GEMME^39^, and mode r=0.672 for the stability predictor DDMut^40^ (Fig. 6b and Extended Data Fig. 4a). The good performance of these non-indel predictors on this large dataset suggests it should be possible to improve indel prediction.

### Accurate indel variant effect prediction using INDELi

Across 171 domains, the average effect of substitutions correlates with deletion tolerance (mode r=0.450) and more weakly with insertion tolerance (mode r=0.253, Fig. 3c and model m1 in Fig 7a-b for comparison). This suggests substitution scores could be repurposed for indel variant effect prediction. Moreover, although the identity of the wildtype aa only poorly predicts deletion and insertion tolerance (model m2, mode r=0.226 and 0.182, respectively, Fig. 7b), a regression model using secondary structure features (strand/helix/loop/termini, length of secondary structure, neighboring structures, and the position of the indel) provides quite good deletion (model m3, mode r=0.538) and insertion (mode r=0.573, Fig. 7b) prediction, when evaluated by leave-one-out cross validation (See Methods, Fig. 7a). This performance is similar to that of the state-of-the-art ESM1b protein language model (Fig. 6a), despite the model containing only 33 parameters.

**Figure 7.**
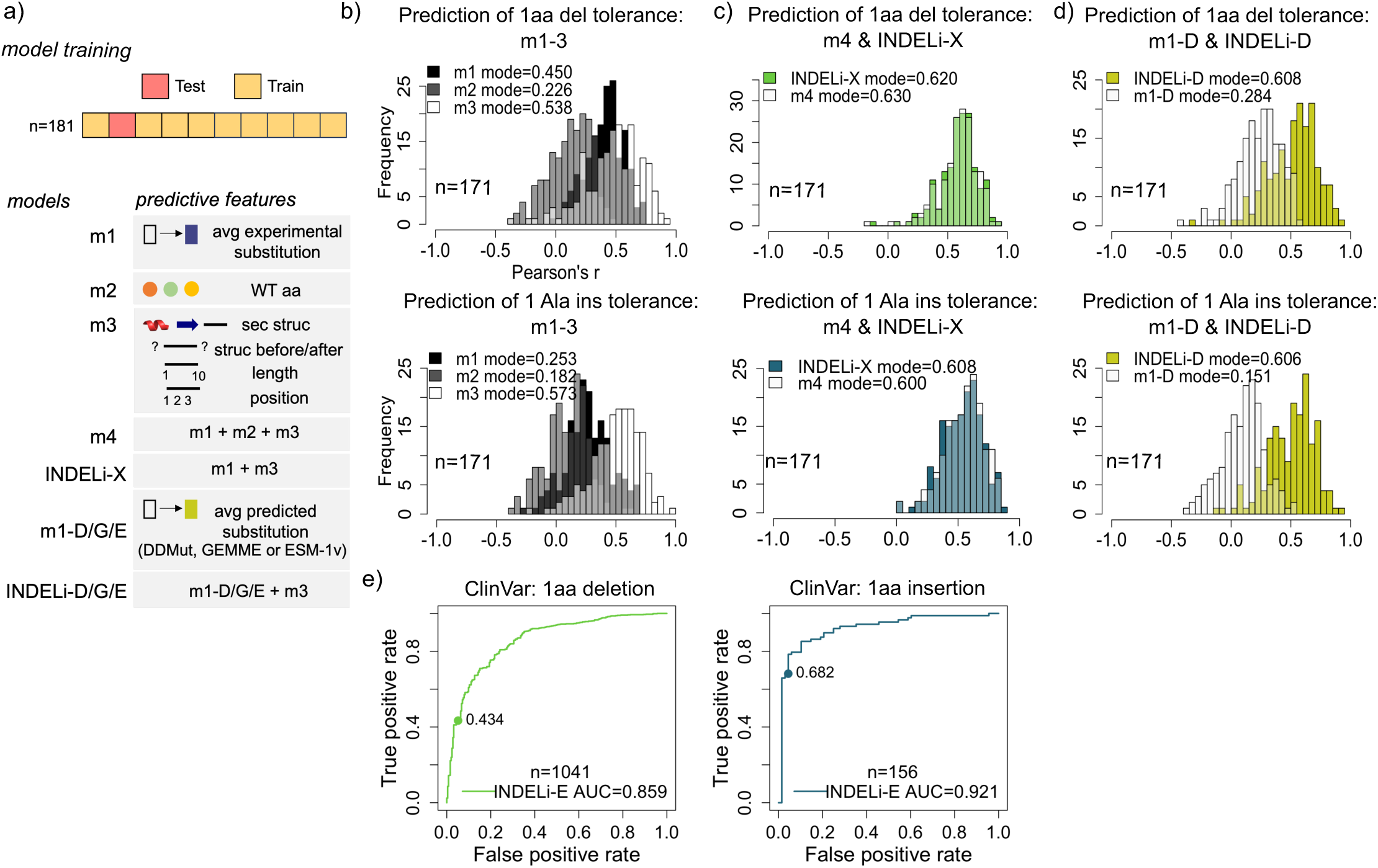
Accurate indel variant effect prediction using INDELi. **a.** Overview of the model training by leave-one-domain out cross-validation, the models used for indel prediction and the predictive features they were trained on (See Methods). **b.** Performance evaluation of models m1-3 for 1aa deletion (upper panel) and 1aa insertion (lower panel) prediction as histograms of Pearson’s correlation between the observed and predicted ddG. Mode r for each model is displayed in the legend. **c.** Performance evaluation of m4 and INDEL-X. **d.** Performance evaluation of m1-D and INDELi-D **e.** Evaluation of the genome-wide prediction of 1aa deletion (left) and insertion (right) effects on stability across the human proteome, using INDELi-E. The performance is evaluated with 1aa indel ClinVar variants (See Methods) using receiver operating characteristics-area under the curve (ROC-AUC). The true-positive rate at 5% of false-positive rate is highlighted on each plot.

We next trained a model that uses both substitution scores and secondary structure features as predictive features. Strikingly, this model has predictive performance similar to state-of-the-art substitution variant effect predictors, with mode r=0.620 for deletions and mode r=0.608 for insertions (Fig. 7c). Moreover, the predictive performance is consistent across domains, with a narrower distribution of correlation coefficients compared to ESM1b (Fig. 7c and Extended Data Fig. 4e, interquartile range from r=0.512 to 0.686 for deletions and r=0.393 to 0.646 for insertions). Incorporating secondary structure information therefore allows experimental substitution scores to be re-purposed to predict the effects of insertions and deletions. We refer to this model as INDELi-X (INDELi using eXperimental substitution scores).

### Accurate genome-wide stability and pathogenic indel prediction using INDELi

To allow genome-wide prediction of indel variant effects, we next tested whether we could replace the experimentally measured substitution scores with computationally predicted ones. We tested the performance of INDELi using three state-of-the-art substitution predictors, DDMut, ESM-1v and GEMME. Similar to experimental measurements, computationally-predicted substitution scores correlate weakly with the effects of deletions (mode r=0.284-0.391) and insertions (mode r=0.151-0.330, Fig. 7d and Extended Data Fig. 4b-c) on protein stability. However, training models that combine these predicted scores with secondary structure features, results in a comparable performance to INDELi-X for both 1aa deletions (mode r=0.608, 0.577 and 0.614 for DDMut, ESM-1v and GEMME) and 1aa insertions (mode r=0.606, 0.587 and 0.553 for DDMut, ESM-1v and GEMME, Fig. 7d and Extended Data Fig. 4b-c). Hence, state-of-the art substitution predictors can be repurposed to accurately predict indel tolerance. We refer to the final models as INDELi-D,-E, and -G for INDELi using DDMut, ESM-1v or GEMME, respectively.

Although the performance of these three models is similar, INDELi-E has the major advantage that it uses ESM-1v substitution predictions, which can be made genome-wide. We therefore used INDELi-E to predict the effects of 1aa insertions and deletions on protein stability across the human proteome. The resulting dataset consists of >14.7 million predictions that we have made available as a resource to evaluate the effects of indels in all human proteins as Extended Data file 8.

INDELi-E is trained to predict the impact of indels on protein stability. Reduced stability is only one of the possible molecular mechanisms by which variants can alter protein function and cause disease. We were therefore interested to test how well a method only trained to predict stability changes could identify insertions and deletions known to cause human diseases. Evaluating INDELi’s predictions on human proteins^41^ suggest that stability changes are a major component of indel effects on function (Extended Data Fig. 4d).

Strikingly, despite only being trained on *in vitro* protein stability, INDELi-E provides very good classification of human pathogenic variants with a receiver operating characteristic curve-area under the curve (ROC-AUC) = 0.859 for 788 pathogenic 1 aa deletions vs. 253 benign and ROC-AUC= 0.921 for 88 pathogenic 1aa insertions vs. 68 benign reported in ClinVar for all human proteins (Fig. 7e). Importantly for clinical applications, INDELi-E also has high sensitivity when making predictions with a low false positive rate (high specificity): for example at a 5% false positive rate, the sensitivity (true positive rate) for identifying pathogenic deletions and insertions is 43.4% and 68.2%, respectively (Fig. 7e). Thus, despite only being trained using high-throughput measurements of the effects of indels on the *in vitro* fold stability of small, mostly microbial, protein domains, INDELi-E provides very good prediction of pathogenic variants in full length multi-domain and structurally diverse human proteins. This excellent performance is consistent with fold stability being the major mechanism by which aa indels cause human genetic diseases.

### Insertions generate gain-of-function molecular phenotypes

Our results for indels and previous analyses for substitutions^42–46^ suggest that destabilization is the most frequent mechanism by which genetic variants in proteins cause human diseases. However, proteins have additional molecular activities beyond folding and an important subset of disease variants have gain-of-function molecular mechanisms^47^. We therefore also quantified the effects of substitutions, insertions and deletions on a biophysical activity beyond fold stability, focussing on protein binding, which is a molecular function important for nearly all proteins. Using a highly-validated quantitative protein-protein interaction selection assay^27,28,48^, (Fig. 8a), we quantified the effects of all aa substitutions and diverse insertions and deletions on the binding of GRB2-SH3 and PSD95-PDZ3 to their respective ligands, GRB2-associated binding protein 2 (GAB2) and CRIPT (Fig. 8c).

**Figure 8.**
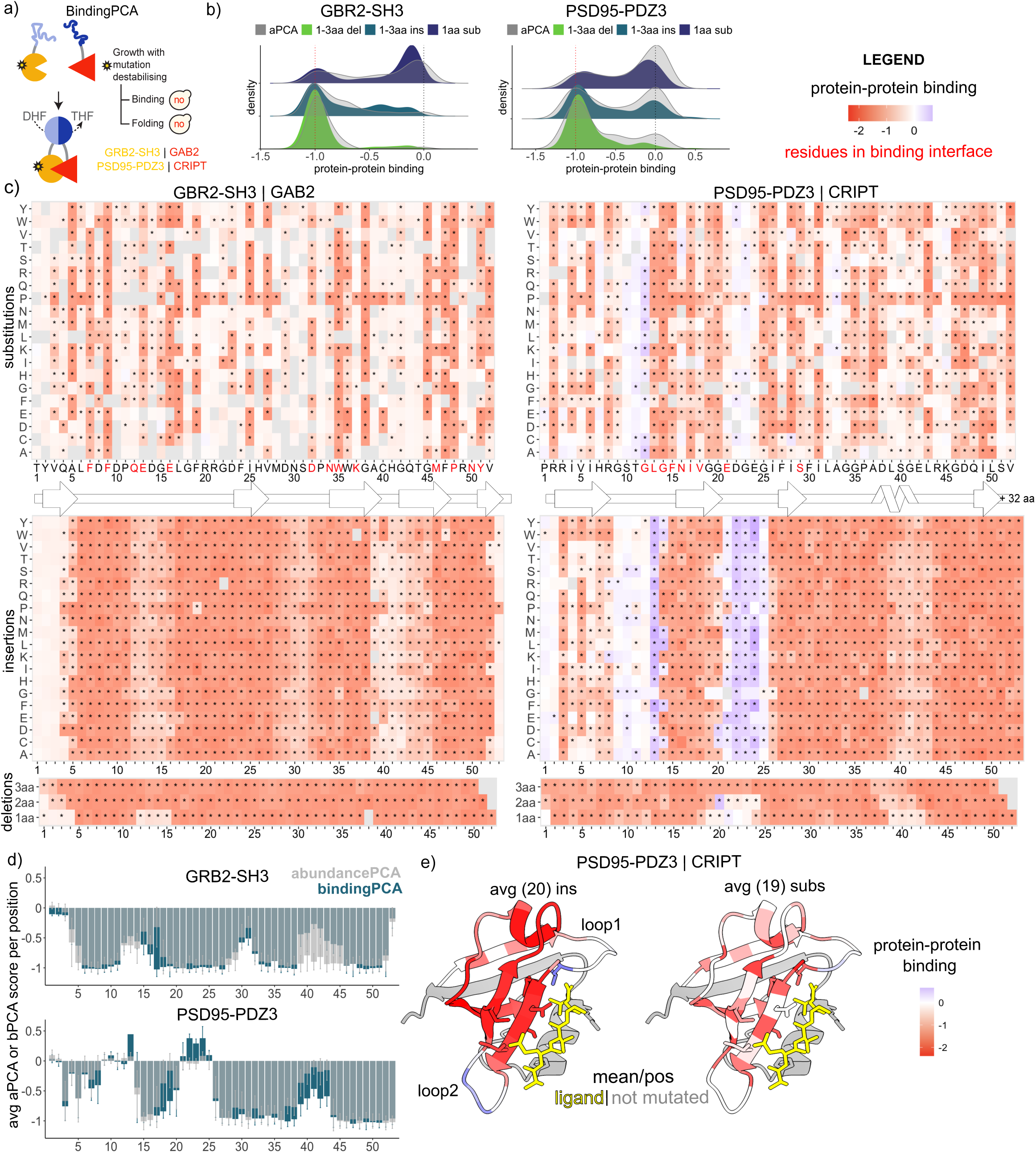
Insertions generate gain-of-function molecular phenotypes. **a.** Overview of the bPCA selections; no: yeast growth defect; DHF: dihydrofolate; THF: tetrahydrofolate**. b.** Density distributions of 1-3 aa deletion (green), 1-3 aa insertion (blue) and single substitution variants (purple); grey: aPCA data. **c**. Heatmaps of 19 single aa substitutions, 20 aa insertions and 1-3aa deletion effects on protein-protein binding. Changes in binding significant from the weighted mean of the synonymous variants are marked with “*” (Bonferroni multiple testing correction of z-stats in a two-tailed test); y-axis: identity or length of mutation; x-axis: mutated position- and the aa sequence of the domain. We show the 2-dimensional structural representation of the domains (SSDraw^63^) under each heatmap. For PSD95-PDZ3 and GRB2-SH3, we mutated the first 52 residues, leaving, respectively, 32 and 4aa in the C-terminal as wild type. **d.** Mean insertion scores/residue for bPCA (blue) and aPCA (grey); error bars: standard deviation of aPCA scores/residue. **e.** Structure of PSD95-PDZ3 interacting with its ligand CRIPT (adapted from pdb 1BE9) coloured by the mean bPCA score of 20 aa insertions/residue (right) and mean of 19 aa substitutions/residue (left). Visualised using ChimeraX^67^.

In both proteins, many insertions and deletions strongly reduce binding to their ligands, consistent with their effects on fold stability (Fig. 8b and Extended Data Fig. 5c). Most strikingly, however, a large number of 1aa insertions in PSD95-PDZ3 have gain-of-function phenotypes, strongly increasing its binding to the CRIPT ligand (Fig. 8c-d). These insertions are nearly all in loops 1 and 2 of the protein (Fig. 8c and e). A subset of multi-aa insertions also increase binding (Extended Data Fig. 5b). Insertions in these loops have a much stronger gain-of-function molecular phenotypes than substitutions (Fig. 8e). Loop 1 constitutes the carboxylate-binding loop, directly involved in peptide recognition and forming the binding site groove for CRIPT together with strand 2 and helix 2^49^. The second loop connects strand 2 and 3, which both contain ligand-contacting residues^49^. The insertions in these loops that increase binding have only a mild effect on the abundance of PSD95-PDZ3 (Fig. 8d)

Thus, although insertions and deletions - like substitutions - primarily have loss-of-function molecular phenotypes, insertions in two loops of PSD95-PDZ3 cause strong gain-of-function phenotypes.

## Discussion

Here we have used deep indel mutagenesis to quantify, understand and learn how to predict the effects of insertions and deletions on protein fold stability. Indels and substitutions are fundamentally different changes to proteins, and this is reflected in their different patterns of tolerance. Our data show that tolerance varies both among different proteins and different regions of the same protein, as does the relative tolerance to insertion and deletion.

Despite this apparent complexity, we have also shown that it is possible to accurately predict the effects on indels on protein stability using relatively simple models that combine experimentally-measured or computationally-predicted substitution scores with information about the secondary structure of a protein. State-of-the-art substitution variant effect predictors can thus be augmented and repurposed as good indel predictors and we have made human genome-wide predictions using one of these models, INDELi-E, available as a resource (Extended Data file 8). Strikingly INDELi-E performs very well as a predictor of pathogenic clinical insertions and deletions, even though it was only trained on the effects of indels on the *in vitro* stability of small, mostly microbial, protein domains. Together with previous work on substitutions^42–46^, this suggests that fold destabilization is the most important mechanism by which aa variants cause human diseases.

Finally, while it has been demonstrated that novel protein functions can be achieved through domain insertions^50–53^ and loop remodelling^54–58^, there has been little systematic evaluation of such effects for short indels^18,52^. Our systematic analysis and previous work^59–61^, illustrate how indels can be a source of gain-of-function phenotypes, which are particularly challenging variants to identify and predict across diseases and are important for evolution^47^.

The expansion of deep indel mutagenesis to additional proteins and to additional molecular and cellular phenotypes is an important goal for future work. Large experimental indel mutagenesis will provide training and evaluation data for computational models for different protein properties and reference atlases to guide the interpretation of clinical variants. Large-scale indel mutagenesis and models to predict the effects of indels are, moreover, likely to be important for protein engineering, enabling more focused sampling of sequence space for directed evolution studies and providing access to gain-of-function and change-of-function phenotypes that are difficult to access through substitutions alone.

## Materials and methods

### Deep Indel mutagenesis library design

Protein domains to mutagenize were chosen to satisfy the following four criteria: (1) structural diversity with the domains covering the major structural classes of all-alpha, all-beta and mixed alpha and beta domains; (2) short length to allow synthesis in a 250nt oligonucleotide pool; (3) the availability of reference *in vitro* ddG folding energy measurements for substitutions in the ProTherm; and (4) good correlation between aPCA growth rates and *in vitro* ddGs for substitutions in a very large substitution mutagenesis screen (Figure 1b and Beltran et al. manuscript in preparation). We designed indel libraries and performed selections for 13 domains but observed weak selection for 4 resulting in high measurement error and weak separation of nonsense and synonymous variants, likely due to insufficient growth time of the pool. We excluded these four domains from the analyses.

We designed the mutational libraries using sequences listed in Extended Data Table 2 as the wildtype templates. For GRB2-SH3 and PSD95-PDZ3 we designed i) 1-3 aa neighboring deletions ii) 1-3 aa neighboring CCC insertions iii) out-of-frame 3nt deletions resulting in delSubs iv) 15x synonymous mutations prioritising >1nt changes v) all possible 19 substitutions at each position using most abundant yeast-codons (https://www.kazusa.or.jp/codon/cgi-bin/showcodon.cgi?species=4932) and prioritising >1nt changes in the substituted codons vi) all possible 20 aa insertions at each position vii) 1-3 aa repeats of each aa in the template sequence viii) 2aa insertions of Cys and viv) 2aa insertions of Ala. For PSD95-PDZ3 and GRB2-SH3, we mutated the first 52 residues, leaving, respectively, 32 and 4aa in the C-terminal as wild type. For the remaining 7 protein domains, we included i-iv into the mutational design. The 5’ and 3’ prime adapters were added to each protein for amplification and cloning purposes and adjusted to each wild type (wt) template length, with final adapters ranging from 20-42nt. The library was ordered at Twist Biosciences as a pooled oligo library with final lengths of single stranded oligos ranging from 149-250nt. The Twist pool was resuspended in water to the concentration of 1 ng/uL and the 3 sub pools 1) GRB2-SH3 2) PSD95-PDZ3 and 3) 7 domains were separated by PCR amplification of 14 cycles, using primers listed in Extended Data Table 3 and using 10ng of the Twist pool as template in each reaction. The 3 library pools were purified by an ExoSAPII (NEB) reaction to remove single-stranded DNA and further by column purification (MiniElute Gel Extraction Kit, QIAGEN).

### Variant library construction

We used generic abundance- and bindingPCA plasmids^27,28^ to construct the 3 sub libraries. GRB2-SH3 and PSD95-PDZ3 libraries were cloned into their respective aPCA plasmids pGJJ046 and pGJJ068 (N-terminal DHFR tag^27^) using Gibson assembly. The 7 domain library was cloned into the pGJJ162 plasmid (C-terminal DHFR tag^28^). The backbones for the Gibson reaction for GRB2-SH3 and PSD95-PDZ3 library assembly (aPCA plasmids) were first linearized using primers listed in Extended Data Table 3 and next treated with Dpn1 (NEB) restriction enzyme to remove the circular plasmid template. Next the correct sizes of the linearized backbones were isolated using gel electrophoresis and later purified using QIAquick Gel Extraction Kit (QIAGEN). For the library of 7 domains, the pGJJ162 backbone was linearised by digestion with restriction enzymes NheI and HindIII (NEB) for the Gibson reaction. We used 100ng of the linearized vectors and 9.4ng, 9.28ng and 11.23ng of the purified double-stranded libraries (GRB2-SH3, PSD95-PDZ3 and 7 domains) for each gibson reaction of 20 uL. The gibson reaction (in-house prepared enzyme mix) was incubated at 50C for one hour, then desalted by dialysis using membrane filters for 1 hour, concentrated to 3X by SpeedVac (Thermo Scientific) and transformed into NEB 10β High-efficiency Electrocompetent E. coli cells according to the manufacturer’s protocol. Cells were allowed to recover in SOC medium (NEB 10β Stable Outgrowth Medium) for 1 hour and later transferred to 100 mL of LB medium with ampicillin overnight. 100 mL of each saturated E. coli culture were harvested next morning to extract the plasmid library using the QIAfilter Plasmid Midi Kit (QIAGEN). Finally, after verifying correct assembly by sanger sequencing (Eurofins), the GRB2-SH3 and PSD95-PDZ3 libraries were digested out of the aPCA plasmids using NheI and HindIII for re-assembly of the bPCA library. The bPCA library was assembled overnight by temperature-cycle ligation using T4 ligase (New England Biolabs) according to the manufacturer’s protocol, 67 fmol of backbone and 200 fmol of insert in a 33.3 uL reaction. As backbones for GRB2-SH3 and PSD95-PDZ3 library inserts, we used the pGJJ034 and pGJJ072 plasmids with GAB2 and CRIPT ligands fused N-terminally to DHFR12, as described in our previous study^27^. The ligation was desalted by dialysis, concentrated 3X, transformed into NEB 10β High-efficiency Electrocompetent E. coli cells, and purified from E. coli using the QIAfilter Plasmid Midi Kit as described above.The coverage of variants after each transformation reaction was estimated to be >20x.

### AbundancePCA and BindingPCA selections

aPCA and bPCA methotrexate selection were performed as described in our previous studies^27,28^. The high-efficiency yeast transformation protocol was scaled to 100 mL based on the targeted number of transformants of each library and each biological replicate was transformed into cells grown from independent colonies of BY4741 yeast strain (https://www.yeastgenome.org/strain/by4741). In short, for each of the selection assays, 3 independent pre-cultures of BY4741 were grown in 20 mL standard YPD media at 30 °C overnight. The next day, the cultures were diluted into 100 mL of pre-warmed YPD at an optical density at 600 nm (OD_600_) of 0.3. The cultures were incubated at 30 °C for 4 h. After incubation, the cells were collected and centrifuged for 5 min at 3,000*g*, washed with sterile water and later with SORB medium (100 mM lithium acetate, 10 mM Tris pH 8.0, 1 mM EDTA, 1 M sorbitol). The cells were resuspended in 4.3 ml of SORB and incubated at room temperature for 30 min. After incubation, 87.5 μl of 10 mg ml^−1^ boiled salmon sperm DNA (Agilent Genomics) was added to each tube with 1.75 μg of plasmid library. After mixing, 17.5 ml of plate mixture was added to each tube to be incubated at room temperature for a further 30 min. DMSO (1.75 ml) was added to each tube and the cells were next heat shocked at 42 °C for 20 min (inverting tubes from time to time to ensure homogeneous heat transfer). After heat shock, cells were centrifuged and resuspended in ∼25 ml of recovery medium and allowed to recover for 1 h at 30 °C. Next, cells were centrifuged, washed with SC-URA medium and resuspended in 100 mL SC-URA. After homogenization by stirring, 10 μl were plated on SC-URA petri dishes and incubated for ∼48 h at 30 °C to quantify the transformation efficiency. The independent liquid cultures were grown at 30 °C for ∼48 h until saturation. The number of yeast transformants obtained in each library assay covered each variant at least 100x.

In total we completed 5 methotrexate selection assays of 3 replicates each, 3 aPCA for GRB2-SH3, PSD95-PDZ3 and the 7-domain library, and 2 bPCA for GRB2-SH3 and PSD95-PDZ3 libraries. The pre-selection medium used was SC-URA/ADE and the selection medium was SC-URA/ADE + 200ug/mL Methotrexate (BioShop Canada Inc., Canada). The selections were performed directly following the big-scale yeast transformation. In short, ∼48 hours after the transformation, cultures were inoculated in the pre-selection media at starting OD600=0.05, taking cells from the saturated cultures post-transformation. Cells were grown for 5 generations at 30 °C under constant agitation at 200 rpm until reaching OD600 ∼1.6. We refer to this step as the library “input”. Next, the selection cultures, referred to as “output”, were inoculated from the “input” cultures at OD600=0.05 and into the selection medium. Cells were collected by centrifugation at 3,000 g and washed with sterile water once to remove the pre-selection media prior to inoculation. The remaining ∼95mL of the “input” cultures were washed with sterile water twice and flash frozen for DNA extraction and sequencing. The “output” cultures were grown for 5 generation at 30 °C under constant agitation at 200 rpm until reaching OD600 ∼1.6. Next, the “output” was collected by centrifugation at 3,000 g and washed twice with sterile water before flash-freeze and storage at –20 °C prior to DNA extraction and sequencing. Harvested input and output cells were pelleted, washed with water and stored at -20C until the DNA extraction step.

### DNA extractions and plasmid quantification

The DNA extraction protocol was used as described in our previous study^27,28^. We extracted DNA from a 50mL harvested selection input and output cultures at OD600nm∼1.6. In short, cell pellets (one for each experiment input or output replicate) were resuspended in 1 ml of DNA extraction buffer, frozen by dry ice-ethanol bath and incubated at 62 °C water bath twice. Next, 1 ml of phenol:chloroform:isoamyl alcohol 25:24:1 (equilibrated in 10 mM Tris-HCl, 1 mM EDTA, pH 8) was added, together with 1 g of acid-washed glass beads (Sigma Aldrich) and the samples were vortexed for 10 min to lyse the cells. Samples were centrifuged at room temperature for 30 min at 4,000 rpm and the aqueous phase was transferred to new tubes. This step was repeated twice. Three molar sodium acetate (0.1 ml) and 2.2 ml of pre-chilled absolute ethanol were added to the aqueous phase. The samples were gently mixed and incubated at −20 °C for 30 min. Next, they were centrifuged for 30 min at full speed at 4 °C to precipitate the DNA. Ethanol was removed and the DNA pellet was allowed to dry overnight at room temperature. DNA pellets were resuspended in 0.6 ml TE and treated with 5 μl of RNaseA (10 mg ml^−1^, Thermo Scientific) for 30 min at 37 °C. To desalt and concentrate the DNA solutions, QIAEX II Gel Extraction Kit was used (20 µl of QIAEX II beads). The samples were washed twice with PE buffer and eluted twice by 75 µl of 10 mM Tris-HCI buffer, pH 8.5 and then the two elutions were combined. Plasmid concentrations in the total DNA extract (that also contained yeast genomic DNA) were quantified by qPCR using the oligo pair 6 (Extended Data Table 3), that binds to the ori region of the plasmids.

### Sequencing library preparation

Sequencing library preparation was done as described in our previous study^27,28^. We performed 2 consecutive PCR reactions for each sample. The first PCR (PCR1) is used to amplify the amplicons for sequencing, to add a part of the illumina sequencing adaptors to the amplicon and to increase nucleotide complexity for the sequencing reaction by introducing frame-shift bases between the adapters and the sequencing region of interest. PCR1 frame-shifting (fs) oligos for each of the sub libraries are listed in Extended Data Table 3. The second PCR (PCR2) is used to add the remainder of the illumina adaptor and to add demulitplexing indexes. All samples, except the GRB2-SH3 bPCA, were dual-indexed using differing barcode indexes both for the forward (5’ P5 Illumina adapter) and reverse oligos (3’ P7 Illumina adapter). The GRB2-SH3 bPCA library was single-indexed using a constant forward oligo (3’ P7 Illumina adapter) and alternating reverse oligos (3’ P7 Illumina adapter). The demulitplexing primers used for PCR2 are listed in Extended Data Table 4. The amplicon library pools were isolated based on size by gel electrophoresis using a 2% agarose gel and then purified using QIAEX II Gel Extraction Kit (QIAGEN) and using 30uL of QIAEX II beads for each sample. The purified amplicons were subjected to Illumina 150bp paired-end NextSeq 2000 sequencing at the CRG Genomics Core Facility.

### Sequence data processing

FastQ files from paired-end sequencing of all aPCA and bPCA experiments were processed with DiMSum^62^ v1.2.11 using default settings with minor adjustments: https://github.com/lehner-lab/DiMSum. Due to low read coverage, following samples were re-sequenced: GRB2-SH3 bPCA input replicate 3, output replicate 1 and 2. Outputs from the second run of sequencing were added as technical replicates in the “Experimental Design File” when running DiMSum for the GRB2-SH3 bPCA experiment. Variant counts associated with all samples (output from DiMSum stage 4) were filtered using a “Variant Identity File”, a list of designed sequences to retain only the programmed variants. In addition to the default DiMSum settings, including a minimum of 10 bp alignment between the pair-end reads, we used the argument “indels” to list the length of expected variants to be retained. FastQ files were processed with DiMSum separately for the GRB2-SH3 and PSD95-PDZ3 aPCA and bPCA samples and in bulk for the remaining 7 domains. For the 7 domains, we also supplied the Synonym Sequence File listing all wt sequences, to retrieve the synonymous variants for all 7 domains. As most of our variants are indels with nt hamming distance =>3 and the library design for substitutions was optimized for 2-3 nt changes, we didn’t apply any filters for the minimum input count in DiMSum and retained all designed variants. Instead, we filtered the DiMSum output (“…fitness_replicates.RData”) based on the mean replicate input count according to the following criteria: 1 nt hamming distance: >=100 mean count, >=2 nt hamming distance: 10 mean count. The typical count for substitutions and indels was 348 and 253 respectively, determined using density estimation. DiMSum .sh files with used settings, the “Variant Identity Files” and the “Experimental Design Files” for all DiMSum runs are available for download at https://zenodo.org/records/10830272 under “DiMSum.zip.

The DiMSum output “…fitness_replicates.RData” files containing the fitness and fitness error estimates were used for further data analysis. After calling the substitution, indel and synonymous variants for each protein, we next normalised the data for each protein separately. To determine the lower limit for normalisation, we applied the Chernoff mode estimator (using the mlv{modeest} function) to identify the mode of the lower peak in the bimodal distribution of all variant effects.This value was subtracted from the DiMSum “fitness” value resulting in “norm_fitness”. The “norm_fitness” of each variant was then divided by the weighted mean of the synonymous variants, resulting in “scaled_fitness”. For one of the proteins, VIL1-HP, we used the “Vieu” mode estimator of the mlv{modeest} function, due to non-overlapping deleterious modes of insertions and deletions. We normalised the replicate errors by dividing the DiMSum fitness sigma with the weighted mean of synonymous variants. Indel variants with >6 and 0 output counts were added back into the working dataframe from the “…variant_data_merge.RData” file, as DiMSum eliminates everything with 0 counts in the output. These variants were only CKS1 mutants (34 out of 432) and were assigned “scaled_fitness” of 0 as they are thought to be completely deleterious for protein abundance. For all variants, we tested if changes in abundance were significant from the weighted mean of the synonymous variants by calculating z-stats with a two-tailed test and using Bonferroni multiple testing correction. The highly deleterious CKS1 variants with 0 counts in the DiMSum output were not marked as significant changes. The density plots illustrating the distributions of normalised abundance scores across different domains and mutation types are available in Extended Data Fig. 1c-d and Extended Data Fig. 2a.

The threshold for considering a mutation deleterious is calculated using kernel density estimation, which we use to define the mode of the peak representing completely deleterious mutants. Next, the upper and lower bounds of this interval are defined using a standard deviation multiplier (in this case, 1 standard deviation), with the upper bound serving as our threshold for deleteriousness.

### Secondary Structure representation using SSDraw

We used SSDraw^63^ for secondary structure representation of the mutated protein domains. The software was installed from https://github.com/ncbi/SSDraw following the provided instructions. The pdb identifiers used for the 9 domains were: 1QQV (VIL1-HP), 2KZG (FBP11-FF1), 1MJC (CSPA-CSD), 1CSP (CSPB-CSD), 2VWF (GRB2-SH3), 1BE9 (PSD95-PDZ3), 1CIQ (CI2A-PIN1), 2HBA (BL17-NTL9) and 1BUH (CKS1). Prior to analysis, the pdb files were downloaded and trimmed corresponding to protein lengths used in the experimental assays (Extended Data Table 2).

### In vitro fold stability data

*In vitro* fold stability data from Tsuboyama et al.^25^ was downloaded from https://zenodo.org/record/7992926. The downloaded files used for further analysis were “Tsuboyama2023_Dataset2_Dataset3_20230416.csv” containing the inferred deltaG and ddG scores for all substitution and indel variants and the “AlphaFold_model_PDBs “ folder containing the pdb files for all assayed domains. For our analysis, we inverted the sign of the inferred “ddG_ML” by multiplying it with -1. Additionally, we filtered the original data frame, retaining only the natural protein domains and domains with scores for all mutation types (19 substitutions, 1aa deletions and 1 Ala insertions) resulting in a final selection of 181 protein domains, spanning over 6,288 positions. When reporting correlations, we consider only protein domains with data for >9 positions (n=171).

### Secondary structure features

Secondary structure assignments were made using STRIDE^64^ using the stride{bio3d} R function. The pdb files used for the structural annotation were those provided for the Tsuboyama et al. dataset at https://zenodo.org/record/7992926. The secondary structure alignment was simplified from 6 categories (“AlphaHelix,” “310Helix,” “Strand,” “Turn,” “Coil,” “Bridge”) by merging annotations for “Turn,” “Coil,” and “Bridge” into a single category called “loop”. Furthermore, we annotated the N- and C-termini as the sequence of amino acids prior to or immediately after the first residue of the first/last secondary structure element (“AlphaHelix”, “310Helix” and “Strand”). Next, we realigned every secondary structure element using the relative solvent accessibility score calculated using PyMol^65^ v.2.3.5. For each structure element (“AlphaHelix”, “310Helix”, “Strand” and “loop”) we first found the median position based on the element length and set the position 0 to the most buried residue (based on the rSASA score) +1/-1 from the median length. Residues towards the N-terminal of the secondary structure element were annotated in negative, descending order (-1, -2, -3 etc) while the residues towards the C-terminal were annotated in ascending order (1, 2, 3 etc). For the N- and C-termini, we annotated the first position immediately prior to or after the secondary structure as position -1/+1. For N-termini we annotated the rest of the positions in negative, descending order while for the C-termini the positions were annotated in an ascending order.

### Variant effect prediction

CADD^34^ was run on the human subset of the domains from Tsuboyama et al.^25^. CADD is a supervised machine-learning model, trained on the binary distinction between simulated *de novo* variants and variants in the human population. Additionally, CADD integrates protein structure, evolutionary constraints and functional predictions into the model. CADD is not trained on genomic variants for which pathogenic or benign status is known but rather it assumes that variants that have arisen and persisted since the last human-ape ancestor are most likely benign, while simulated variants that are not present in the human population will be most likely pathogenic. This is one of the key advantages of the model as it’s trained on a much bigger training data set than if using clinical annotations. The chromosomal coordinates of each aa sequence were manually annotated using the UCSC Genome Browser^66^ (https://genome.ucsc.edu/index.html) and the VCF files containing all substitutions and single indels coordinates together with the reference- and alternative sequences submitted through the CADD web interface at https://cadd.gs.washington.edu/score.

PROVEAN^35^ predictions were run locally using version 1.1.5, available for download at: https://www.jcvi.org/research/provean#downloads. PROVEAN is an unsupervised model, trained on a clustering of BLAST hits for a query sequence. A delta alignment score is computed for each sequence hit, and the scores are averaged within the clusters of hits. PROVEAN aligns homologous sequences from different organisms to identify the conservation of each position within the sequence to compute the likeliness of a variation from sequence to be likely benign or pathogenic. For the PROVEAN input we used the wildtype aa sequences from the Tsuboyama et. al. dataset from which we encoded all possible substitutions and single indels. The correlation, reflecting the agreement between predicted substitution and indel scores from CADD and PROVEAN, can be found under Extended Data Fig. 4f-g.

ESM1v^38^ (https://github.com/facebookresearch/esm), a large protein language model, was run with minor modifications to the code (predict.py) to allow execution on a CPU with a pytorch installation without CUDA support.

DDMut^40^, a stability predictor trained on ProTherm, was run using the Application Programming Interface (API) with further instructions available at https://biosig.lab.uq.edu.au/ddmut/api. We encoded all possible substitutions using wildtype aa-sequences from the Tsuboyama et. al. dataset. The pdb files used for the DDMut submission were those provided for the Tsuboyama et al. dataset at https://zenodo.org/record/7992926.

ESM1b^37^, a large protein language model, was downloaded following the instructions at: https://github.com/ntranoslab/esm-variants and run locally to predict all possible substitutions, single Ala insertions and single deletions using the wildtype aa-sequences from the Tsuboyama et. al. dataset.

### Model design

For the deletion and insertion prediction models we used multiple linear regression with lasso regularisation without interaction terms, which allowed us to fit a simple linear model to the data while encouraging sparsity of the predictive features by shrinking some regression coefficients to zero. The predictive features (dummy-) encoded for the models 1-4 and INDELi were: “resid”, “simple_struc”, “structure_before”, “structure_after”, “align_to_centre”, “align_to_centre_termini”, “length” in addition to continuous, numeric features “ddG_ML_subs”, “ddMut”, “esm1v” and “GEMME”. The “resid” had 20 levels which described the wildtype aa of the deletion position or the wildtype position before the inserted aa. The 5 levels of secondary structure elements (“AlphaHelix”, “Strand”, “310Helix”, “loop” and “termini”) were encoded by “simple_struc”. The “structure_before” and “structure_after” encoded the structural element immediately before or after the current element and had 6 levels (“start”/”end”, “ntermini”/”ctermini”, “loop”, “Strand”, “AlphaHelix”, “310Helix”). The re-aligned position of each secondary structure element was encoded by “align_to_centre” and was simplified to 9 levels describing the position 0, the 1st, 2nd and 3rd position prior to or after 0, while the rest of the residues were labelled as “>4+” or “-4<”. The same strategy was applied for simplifying the encoding of termini positions in “align_to_centre_termini”. For the “length” we encoded simplified length of each secondary structure element described by 3 levels, “short”, “medium” and “long”, which was based on the frequency of the the element lengths across the Tsuboyama et al. dataset and adapted to each secondary structure element individually. Helices of lengths 4-8 aa were defined as “short, 9-15 aa as “medium” and >16 aa as long. Strands of lengths 2-4 aa were defined as “short, 5-6 aa as “medium” and >7 aa as long. All 310-helices (lengths 3-6 aa) were classified as “medium”. Loops of 1-3 aa were classified as “short”, 4-5 aa as “medium” and >6 aa as “long”. For N- and C-termini lengths of 1-3 aa were classified as “short” and >4 as “long”. Finally, we used the average ddG of substitutions/residue, “ddG_ML_subs”, as a predictive feature for INDELi-X and “ddMut”, “esm1v” and “GEMME”, as the average predicted ddG of stability for 19 substitutions per residue for INDELi-D, -E and -G. The models were evaluated using leave-one-out cross-validation where the model was trained on all-except-one domain and evaluated on that held-out domain. For each individual model, using R, we first determined the optimal regularisation parameter (lambda) by cross-validation using the cv.glmnet{glmnet} function with lasso penalty (alpha=1). Here we also calculated the best lambda value, representing the optimal regularisation strength that minimises overfitting while maintaining model performance. The best-fitting model was trained on all-except-one domain using the glmnet{glmnet} function with the selected optimal lambda from the previous step. Finally, the model performance was tested on the held-out domain. Therefore, in conclusion, the deletion and insertion predictors were tested 181 times for each domain. The final figures contain evaluations of the models for 171 domains as we filtered for >9 positions with indel/substitution measurements, when calculating the correlation coefficients. The top significant model coefficients for INDELi-X and INDELi-D are available in Extended Data Fig. h-i.

### Genome-wide prediction of indel effects using INDELi-ESMv1

The genome-wide prediction was performed on the AlphaFold predictions of the human proteome, downloaded at: https://alphafold.ebi.ac.uk/download#proteomes-section. Secondary structure assignments were made using STRIDE^64^ using the stride{bio3d} R function as described in the “Secondary structure features” method section using the AlphaFold pdb files. Relative solvent accessibility score was calculated using PyMol^65^ v.2.3.5. For the genome-wide prediction, we excluded proteins that i) were longer than 1024 aa due to ESM-1v^38^ prediction length restriction ii) proteins with only one predicted structure iii) proteins that contained PI-helices (missing in the INDELi training data) and iv) proteins with loop, turn or bridges as the only assigned structure. Secondary structure features were encoded in R, using custom functions “structural_features.R” and “encoded_data_for_predictor.R” as described in the section “Model design”, available at https://github.com/lehner-lab/deep_indel_mutagenesis. Structural features were dummy-encoded for the indel effect prediction. For the genome-wide prediction, we trained the INDELi-ESM-1v model on all 181 protein domains from Tsuobyama et al dataset^25^ as described in the section “Model design”. Overall, we provide 14,750,162 genome-wide predictions for effects of 1aa deletions and insertions on protein stability, across 17,851 human proteins (Extended Data file 8).

### Selection of ClinVar variants

The set of all ClinVar variants was downloaded from the FTP site on 5th of March 2024 (https://ftp.ncbi.nlm.nih.gov/pub/clinvar/vcf_GRCh38/clinvar_20240301.vcf.gz). For the validation of the genome-wide prediction, we isolated all in-frame deletions and insertions of 3nt causing a single aa change. Furthermore, we ran CADD and PROVEAN predictions on the isolated variants as described in the “Variant Effect Prediction” method section. The ClinVar variants were matched to the genome-wide predictions with INDELi-ESM-1v by running a RefSeq to Uniprot mapping on the Uniprot server (https://www.uniprot.org/id-mapping). The final ClinVar benchmark dataset of 1 aa indels contains 4,630 deletions and 800 insertions and is provided as Extended Data file 6 and 7. For 1aa deletions, 253 variants are benign and 788 pathogenic, the rest being variants of unknown significance. For 1aa insertions, we retrieve 68 benign and 88 pathogenic variants.

## Data availability

All DNA sequencing data have been deposited in the Gene Expression Omnibus under the accession number GSE244096. All scaled fitness measurements for i) aPCA, ii) aPCA of selected substitution mutants for the *in vitro* ddG validation, iii) bPCA iv) the processed Tsuboyama et al. data for indels and mean substitutions/residue v) all single substitutions vi) filtered 1aa deletions and vii) insertion ClinVar variants and viii) genome-wide 1aa indel predictions are available as Extended Data files 1-8 respectively.

## Code availability

Source code used to perform all analyses and to reproduce all figures in this work is available at: https://github.com/lehner-lab/deep_indel_mutagenesis. Data required to run the source code is available at: https://zenodo.org/records/10830272. We provide code to run the pre-trained INDELi-E models to predict 1aa indel effects in a protein of interest at: https://github.com/lehner-lab/deep_indel_mutagenesis/tree/main/single_protein_prediction.

## Author contributions

M.T. performed all experiments and analyses except the generation of aPCA substitution data for comparison with *in vitro* ddG measurements and ESM1v predictions that were performed by T.B.. M.T. and B.L. conceived the project, designed analyses, and wrote the manuscript with input from T.B.

## Acknowledgements

This work was funded by a European Research Council (ERC) Advanced (883742) grant, the Spanish Ministry of Science and Innovation (LCF/PR/HR21/52410004, EMBL Partnership, Severo Ochoa Centre of Excellence), the Bettencourt Schueller Foundation, the AXA Research Fund, Agencia de Gestio d’Ajuts Universitaris i de Recerca (AGAUR, 2017 SGR 1322), and the CERCA Program/Generalitat de Catalunya. M.T. was funded by a Spanish Ministry of Science and Innovation Severo-Ochoa fellowship PRE2020-093984-SO). T.B was funded by an EMBO (ALTF 183-2020) and Marie Skłodowska-Curie (101030961) fellowship.

## Figure legends

**Extended Data Figure 1.**
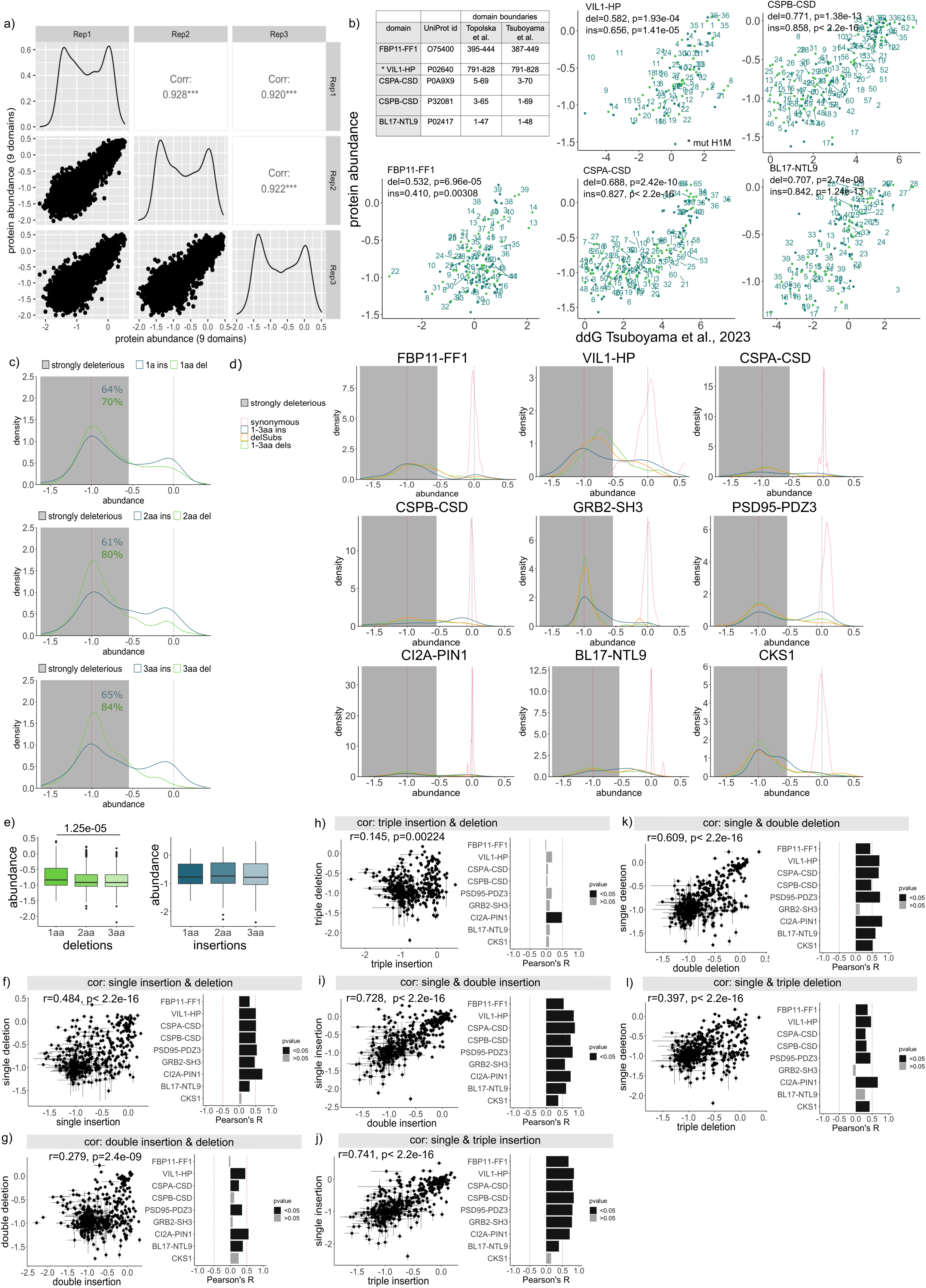
Experimental reproducibility and comparisons of single and multi-aa indels. **a.** Scatter plots showing reproducibility of abundance scores for the 9 domains. **b**. Scatter plots showing correlation of aPCA scores and inferred ddG stability scores from Tsuboyama et al., for indels across the overlapping 5 domains. Pearson’s correlation reported for each domain. The table reports on differences in domain boundaries across the experiments; green: deletions; blue: insertions. **c.** Density distributions of 1-3aa deletions (green) and 1-3aa CCC insertions (blue) aPCA scores for all 9 domains. d. Density distributions of all mutation types for the 9 domains. **e.** Variation in indel deleteriousness across 1-3aa indel lengths. P-values were calculated using Wilcox one-sided t-test with multiple testing adjustment (Bonferroni). **f-l.** Scatter plots of 1-3aa insertion versus deletion aPCA scores (f-h) and single versus multi-aa insertion or deletion aPCA scores (i-j, k-l respectively) across the 9 domains. Results are reported as scatter plots for all 9 domains and bar plots with Pearson’s correlation coefficients for each individual domain.

**Extended Data Figure 2.**
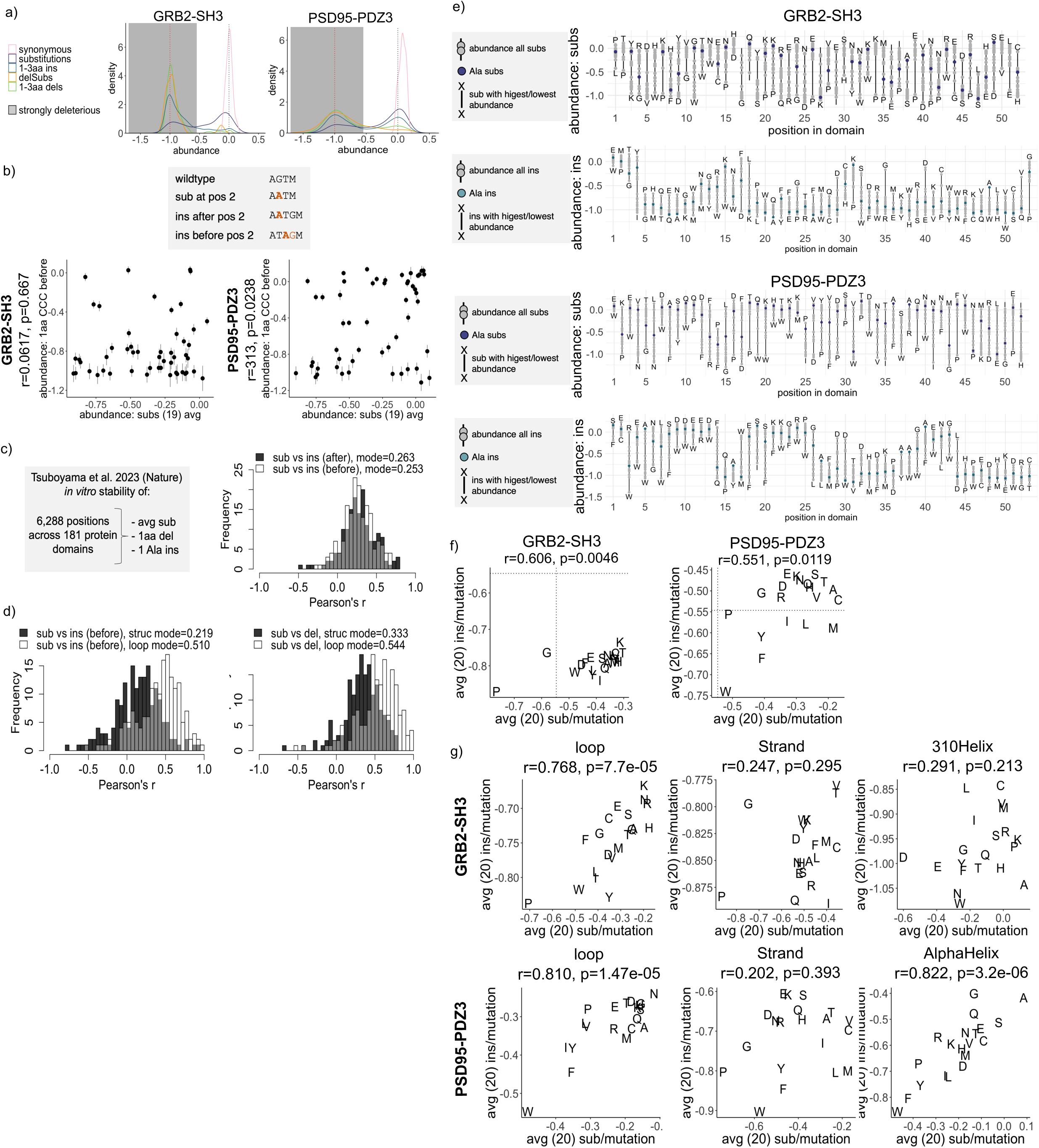
Comparison of substitutions and insertion effects. **a.** Density distributions of all mutation types for GRB2-SH3 and PSD95-PDZ3, including all possible 20 insertions at each position. **b.** Scatter plots of 1aa CCC insertion versus effects of mean substitution/residue with Pearson’s correlations and corresponding p-values. **c.** Overview of the filtered Tsuboyama et al. dataset (See Methods) and histogram of Pearson’s correlations between average substitution per position ddG and 1 Ala insertion before or after the substitution site. **d.** Histograms of Pearson’s correlations between the all mutational types within loop (white) and structured (black) regions for the Tsuobyama et al. dataset. **e.** Domain abundance scores of 19 aa substitutions (upper panel) and 20 aa insertions (lower panel) at each position; purple dot: substitutions to Ala, blue dot: insertions of Ala **f.** Scatter plots of mean insertion and mean substitution scores/aa identity with Pearson’s correlations. The dotted lines mark the threshold of severe deleteriousness for protein domain abundance (See Methods). **g.** Scatter plots of mean insertion and mean substitution scores per aa identity within loops, strands, 310-helices and alpha-helices.

**Extended Data Figure 3.**
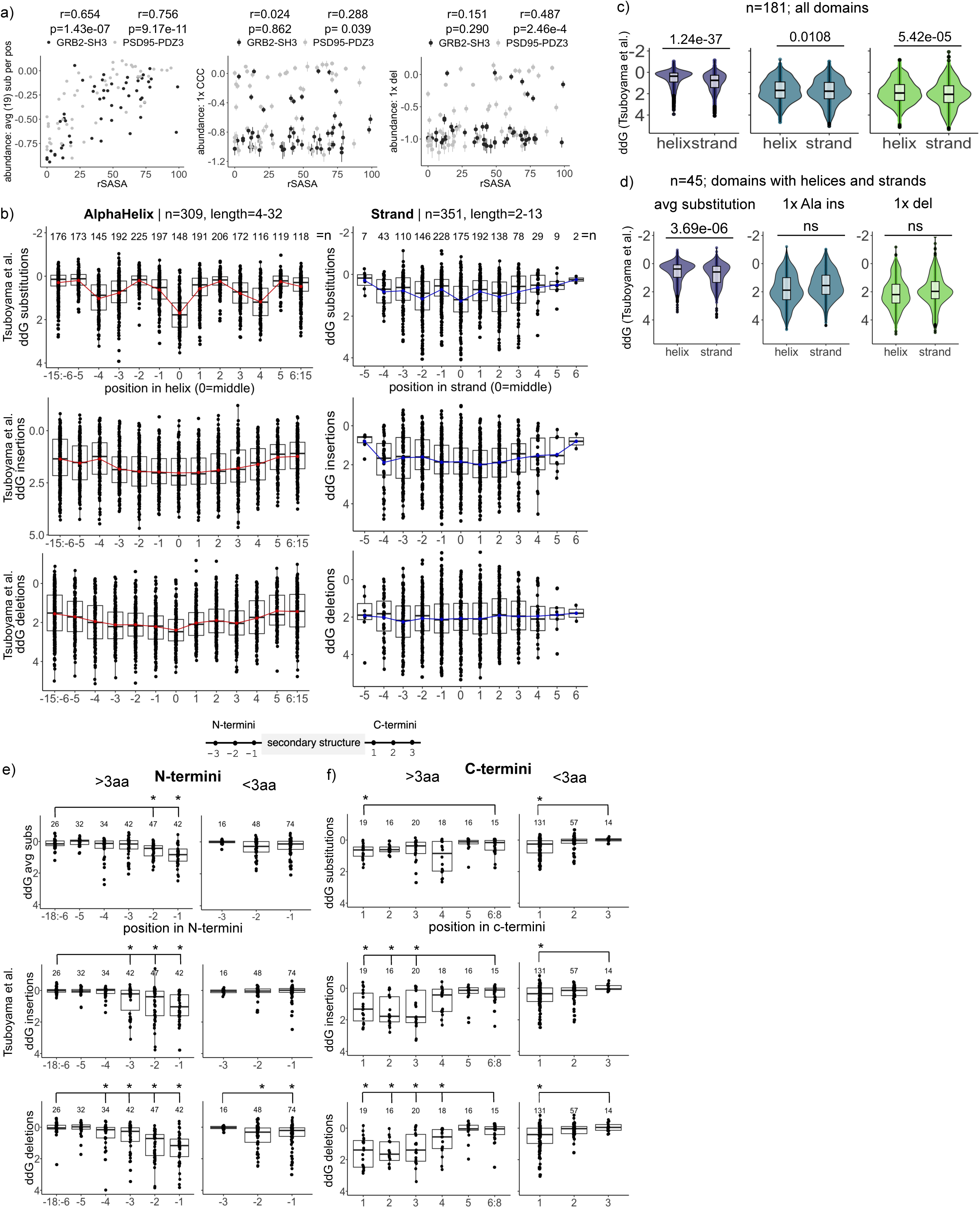
Variability of indel tolerance across secondary structure features. **a.** Scatter plots of protein abundance scores for average substitution/residue, 1aa CCC insertion and 1aa deletions versus the relative solvent accessible area (rSASA) with Pearson’s correlation coefficients. **b.** Patterns of ddG periodicity for substitutions, 1 Ala insertions and 1aa deletions across alpha-helices (red) and strands (blue). Red/blue line represents the mean/position. Position in helix/strand on the x-axis is counted from the middle of the structural element (See Methods). **c.** Violin plots of average substitution, 1 Ala insertion and 1aa deletion ddG between alpha-helices and strands across all 181 domains and **d.** across protein domains that contain both alpha-helices and strands. P-values from Wilcox one-sided t-tests are displayed above the plots. ns: p-value>0.05. **e-f.** Average substitution, 1 Ala insertion and 1aa deletion tolerance across short (1-3aa) and long (=>4aa) N- and C-termini. Numbers of termini/position are indicated above the plots; x-axis: realigned termini positions explained by the legend above the plot. Significant (Wilcox one-sided t-test pvalue < 0.05) changes are denoted by “*”.

**Extended Data Figure 4.**
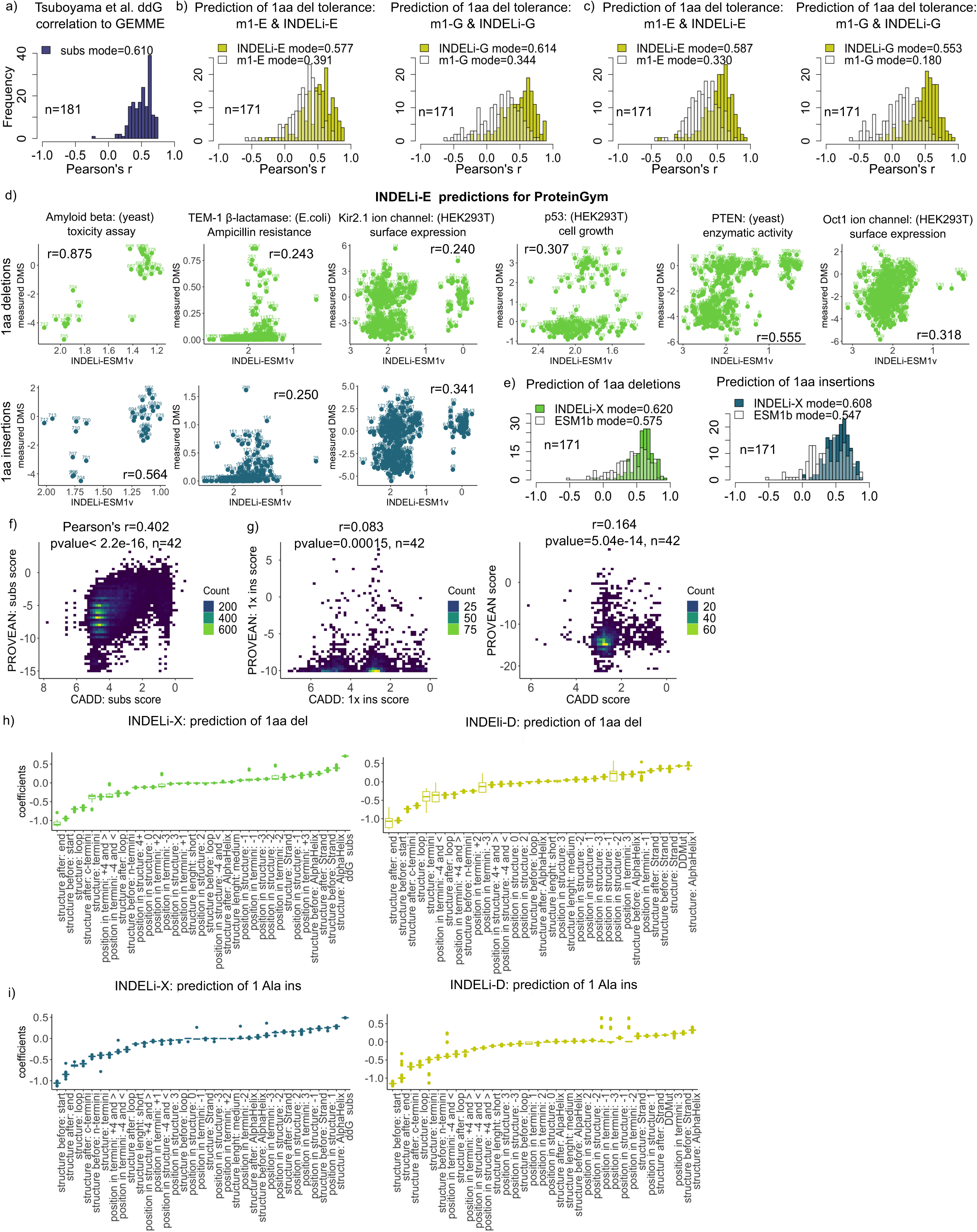
Indel variant effect prediction. **a.** Histograms of Pearson’s correlation coefficients for observed and predicted tolerance scores (>9) of average substitutions for the Tsuboyama et al. dataset using GEMME^39^; n=181. **b-c.** Performance evaluation of models m1-E and -G and INDELi-E and -G for 1aa deletion (b) and 1aa insertion (c) prediction, as histograms of Pearson’s correlation between the observed and predicted ddG, using ESM-1v^38^ and GEMME. Mode r for each model is displayed in the legend. **d.** Scatter plots for observed and predicted 1aa deletion (green) and 1aa insertion (blue) scores across the indel ProteinGym benchmark dataset (https://proteingym.org/), not already included in the leave-one-domain-out cross-validation, using INDELi-E. **e.** Direct comparison of ESM-1b^37^ performance compared to INDELi-X shown as histograms of Pearson’s correlation between the observed and predicted ddG. **f.** Plots of correlations between the predicted PROVEAN and CADD substitution scores. Results are shown as a scatter plot and histogram with the per-domain Pearson’s correlation. **g.** Scatter plots of the correlation between the predicted PROVEAN and CADD insertion and deletion scores and the corresponding Pearson’s correlation. **h-i.** Top significant coefficients from the deletion (f) and insertion (g) predictor models INDELi-X (left) and INDELi-D (right).

**Extended Data Figure 5.**
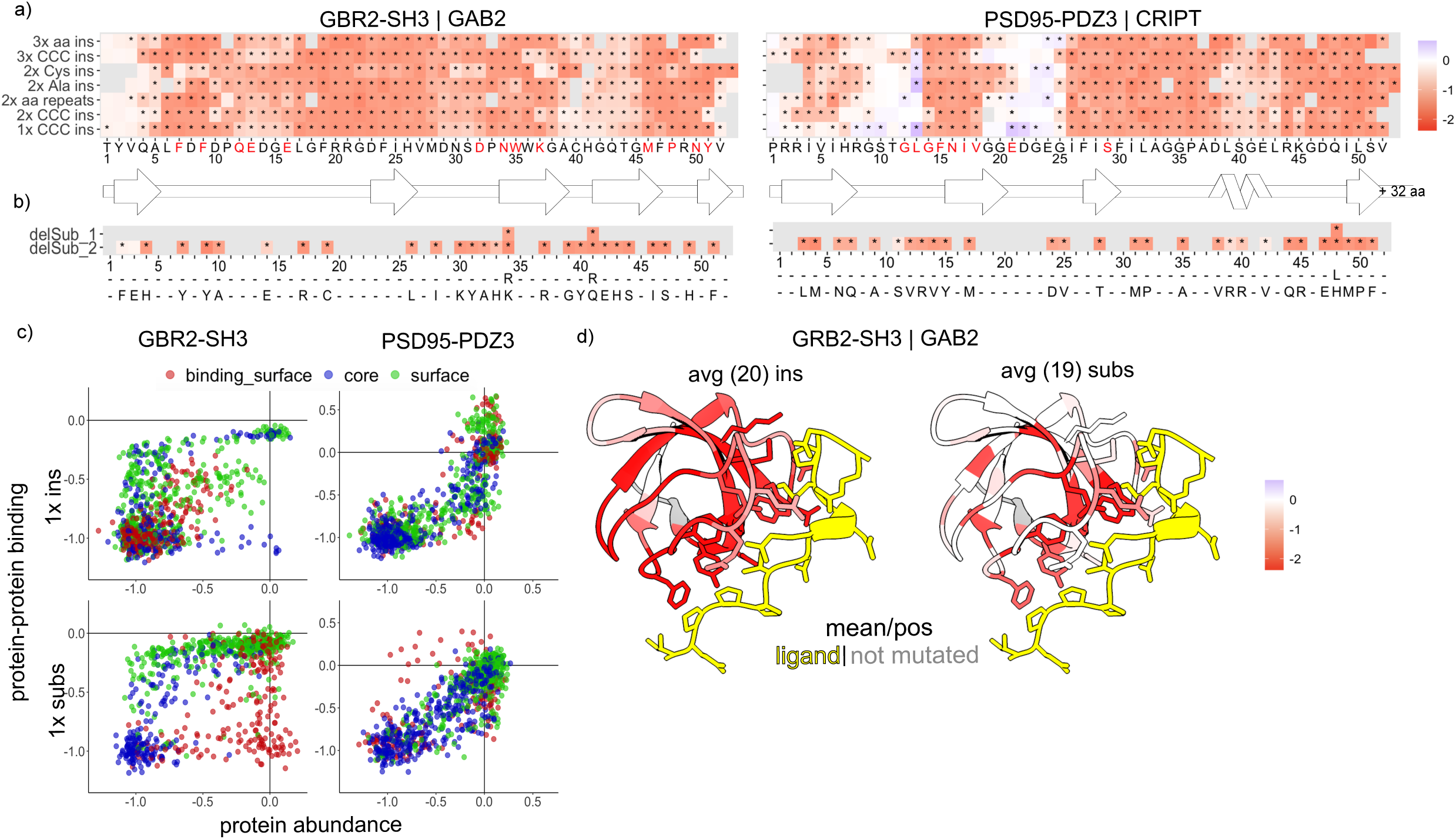
Impact of multi-aa insertions and delSubs on protein interactions. **a-b.** Heatmaps of various types of 1-3aa insertions (a) and delSubs (b) bPCA scores. Changes in binding significant from the weighted mean of the synonymous variants are marked with “*” (Bonferroni multiple testing correction of z-statistic in a two-tailed test); y-axis: type of mutation; x-axis: mutated position and identity of the substitution for delSub. We show the 2-dimensional structural representation of the domains (SSDraw^63^) under each heatmap. For PSD95-PDZ3 and GRB2-SH3, we mutated the first 52 residues, leaving, respectively, 32 and 4aa in the C-terminal as wild type. **c.** Scatter plots of 19 substitution and 20 insertion effects on protein domain abundance and binding; red: residues in binding surface; blue: residues with rSASA<30; green: residues with rSASA>30. **d.** Structure of GRB2-SH3 interacting with its ligand GAB2 (pdb 2VWF) coloured by the mean bPCA score of 20 aa insertions/residue (right) and mean of 19 aa substitutions/residue (left). Visualised using ChimeraX^67^.

