## Extended data tables for "Deep indel mutagenesis reveals the impact of amino acid insertions and deletions on protein stability and function"

Extended Data Table 1. List of Variant effect predictor generated by Atlas of Variant Effects. Available at: https://docs.google.com/spreadsheets/d/17Zsqz-VIS8HQ_Rfl135zRozjyEEcvf9sYdBkxoXwWqc/edit#gid=1328608121

| Predictor | Reports on indels? | Indel scores? | link: |
|---|---|---|---|
| **SIFT - Sorting Intolerant From Tolerant** | yes | Yes/probability score of pathogenicity (0-1) | https://sift.bii.a-star.edu.sg/ |
| **PolyPhen-2 - Polymorphism Phenotyping v2** | no | | http://genetics.bwh.harvard.edu/pph2/ |
| **SNPs&GO** | no | | https://snps.biofold.org/snps-and-go/snps-and-go.html |
| **PROVEAN - PROtein Variant Effect ANalyzer** | yes | yes | https://www.jcvi.org/research/provean#downloads |
| **Condel - Consensus deleteriousness score** | no | | http://bbglab.irbbarcelona.org/fannsdb/ |
| **MutPred2** | yes | Yes/probability score of pathogenicity (0-1) | http://mutpred.mutdb.org/ http://mutpred2.mutdb.org/mutpredindel/ |
| **SNPs&GO3D** | no | | https://snps.biofold.org/snps-and-go/snps-and-go-3d.html |
| **PON-P2 - Pathogenic Or Not Pipeline v2** | no | | http://structure.bmc.lu.se/PON-P2/ |
| **MutationAssessor** | no | | http://mutationassessor.org/r3/ |
| **fathmm - f**unctional **a**nalysis through **H**idden **M**arkov **M**odels | yes | Yes/probability score of pathogenicity (0-1) | http://fathmm.biocompute.org.uk/ http://indels.biocompute.org.uk |
| **CADD - Combined Annotation Dependent Depletion** | yes | yes | https://cadd.gs.washington.edu/ |
| **MutationTaster2021** | yes | classifier | https://www.genecascade.org/MutationTaster2021/ |
| **SuSPect - Disease Susceptibility-based SAV Phenotype Prediction** | no | | http://www.sbg.bio.ic.ac.uk/suspect/index.html |
| **MetaSVM & MetaLR** | no | | https://academic.oup.com/hmg/article/24/8/2125/651446 |
| **SNAP2** | no | | https://rostlab.org/services/snap2web/ |
| **M-CAP - Mendelian Clinically Applicable Pathogenicity score** | no | | http://bejerano.stanford.edu/mcap/ |
| **DEOGEN2** | no | | http://babylone.3bio.ulb.ac.be/MutaFrame/ |
| **Envision** | no | | https://envision.gs.washington.edu/shiny/envision_new/ |
| **FATHMM-XF - FATHMM with an eXtended Feature set** | no | | https://fathmm.biocompute.org.uk/fathmm-xf/ |

| | | | |
|---|---|---|---|
| **DeepSequence** | no | | https://github.com/debbiemarkslab/DeepSequence |
| **ClinPred** | no | | https://www.ncbi.nlm.nih.gov/pmc/articles/PMC6174354/ |
| **EVE - Evolutionary model of Variant Effects** | no | | https://evemodel.org/ |
| **MVP - Missense Variant Pathogenicity prediction** | no | | https://www.nature.com/articles/s41467-020-20847-0 |
| **VARITY** | no | | http://varity.varianteffect.org/ |
| **LRT - Likelihood Ratio Test** | no | | https://www.ncbi.nlm.nih.gov/pmc/articles/PMC2752137/ |
| **EVmutation** | no | | https://marks.hms.harvard.edu/evmutation/downloads.html |
| **LIST_S2** | no | | https://academic.oup.com/nar/article/48/W1/W154/5827198?login=true |
| **MTBAN - Mutation effect prediction using the Temporal convolutional network and the Born-Again Networks** | no | | https://www.nature.com/articles/s41598-021-98693-3 |
| **VESPA - Variant Effect Score Prediction without Alignments** | no | | https://link.springer.com/article/10.1007/s00439-021-02411-y |
| **ESM-1v** | no | | https://github.com/facebookresearch/esm |
| **GEMME - Global Epistatic Model for predicting Mutational Effects** | no | | http://www.lcqb.upmc.fr/GEMME/submit.html |
| **VEST4 - Variant Effect Scoring Tool 4** | yes | Yes/probability score of pathogenicity (0-1) | https://www.cravat.us/CRAVAT/help.jsp |
| **PON-PS - Pathogenic Or Not Pipeline, Severity** | no | | http://structure.bmc.lu.se/PON-PS/ |
| **FATHMM-MKL - FATHMM-Multiple Kernel Learning** | no | | http://fathmm.biocompute.org.uk/fathmmMKL.htm |
| **PrimateAI** | no | | https://github.com/Illumina/PrimateAI |
| **PaPI** | yes | Yes/probability score of pathogenicity (0-1) | http://papi.unipv.it/help.xhtml |
| **DANN** | no | | https://academic.oup.com/bioinformatics/article/31/5/761/2748191?login=true |
| **MetaRNN** | yes | Yes/probability score of pathogenicity (0-1)? | https://genomemedicine.biomedcentral.com/articles/10.1186/s13073-022-01120-z |

| | | | |
|---|---|---|---|
| **BayesDel** | "yes": uses CADD score | | https://onlinelibrary.wiley.com/doi/10.1002/humu.23158 |
| **Sequence UNET** | no | | https://journals.plos.org/plosgenetics/article?id=10.1371/journal.pgen.1008922 |
| **DeepSAV** | no | | http://prodata.swmed.edu/DBSAV/index.html |
| **UNEECON - UNified inferencE of variant Effects and gene CONstraints** | no | | https://journals.plos.org/plosgenetics/article?id=10.1371/journal.pgen.1008922 |
| **CPT - Cross Protein Transfer** | no | | https://genomebiology.biomedcentral.com/articles/10.1186/s13059-023-03024-6 |
| **SAAPpred - Single Amino Acid Polymorphism prediction** | no | | http://www.bioinf.org.uk/mutations/saapdap/ |
| **LASSIE - Linear Allele-Specific Selection InferencE** | no | | https://www.ncbi.nlm.nih.gov/pmc/articles/PMC6673719/ |
| **Rhapsody** | no | | http://rhapsody.csb.pitt.edu/ |
| **COSMIS - COntact Set MISsense tolerance** | no | | https://www.nature.com/articles/s41467-022-30936-x |
| **MISTIC - MISsense deleTeriousness predICtor** | no | | http://lbgi.fr/mistic/ |
| **Tranception** | yes | yes | https://github.com/OATML-Markslab/Tranception |
| **ProtTrans** | no | | https://ieeexplore.ieee.org/document/9477085 |
| **VariPred** | no | | https://www.biorxiv.org/content/10.1101/2023.03.16.532942v1.full.pdf |
| **3Cnet** | no | | https://academic.oup.com/bioinformatics/article/37/24/4626/6322986?login=true |
| **DAMpred - Disease Associated Mutation predictor** | no | | https://zhanggroup.org/DAMpred/ |
| **EvoRator2** | no | | https://www.sciencedirect.com/science/article/pii/S0022283623002401 |
| **PrimateAI-3D** | no | | https://primad.basespace.illumina.com/help |
| **MLVar** | yes | classifier | https://www.nature.com/articles/s41598-022-06547-3 |
| **CAPICE - Consequence-Agnostic prediction of Pathogenicity Interpretation of Clinical Exome variations** | yes | not sure, server is down | https://genomemedicine.biomedcentral.com/articles/10.1186/s13073-020-00775-w |

| | | | |
|---|---|---|---|
| **InMeRF - Individual Meta RF** | no | | https://www.med.nagoya-u.ac.jp/neurogenetics/InMeRF/ |
| **DeMaSk** | no | | https://demask.princeton.edu/about/ |
| **ENTPRISE - sequence ENTropy and PRedIcted protein StructurE for predicting human disease-assiciated amino acid variations.** | no | | https://journals.plos.org/plosone/article?id=10.1371/journal.pone.0150965 |
| **VIPUR - Variant Interpretation and Prediction Using Rosetta** | no | | https://academic.oup.com/nar/article/44/6/2501/2499465?login=true |
| **EA - Evolutionary Action** | no | | https://genome.cshlp.org/content/24/12/2050.long |
| **MOI-Pred - Mode Of Inheritance-Predictor** | no | | https://github.com/rondolab/MOI-Pred/ |
| **LYRUS - Lai Yang Rubenstein Uzan Sarkar** | no | | https://github.com/jiaying2508/LYRUS |
| **Meta-SNP** | no | | https://bmcgenomics.biomedcentral.com/articles/10.1186/1471-2164-14-S3-S2 |
| **UMD-Predictor** | no | | https://umd-predictor.genomnis.com/ |
| **DeepRank-Mut** | no | | https://github.com/DeepRank/DeepRank-Mut |
| **gMVP - graphical Missense Variant Pathogenicity Predictor** | no | | https://github.com/ShenLab/gMVP/ |
| **SNPred** | no | | https://github.com/ArtomovLab/SNPred |
| **AlphaMissense** | no | | https://www.science.org/doi/10.1126/science.adg7492 |

Extended Data Table 2. Amino acid sequence of the wildtype domains and overhangs used for library design.

| **wt_name** | **wt_nt_seq** | **overhang 5'** | **overhang 3'** |
|---|---|---|---|
| psd95-pdz3 | CCGAGGCGAATTGTGATCCACCGGGGCTCCACGGGCCTGGGCTTCAACATCGTGGGTGGCGAGGACGGTGAAGGCATCTTCATCTCCTTTATCCTGGCCGGGGGCCCTGCAGACCTCAGTGGGGAGCTGCGGAAGGGGGACCAGATCCTGTCGGTC | CACTACAAGTACCGTCGACACCGGCTCGGGAGGTGGAGCTAGC | AACGGTGTGGACCTCCGAAATGCTGTGAACGCGCAGATGATC |
| grb2-sh3 | ACATACGTCCAGGCCCTCTTTGACTTTGATCCCCAGGAGGATGGAGAGCTGGGCTTCCGCCGGGGAGATTTTATCCATGTCATGGATAACTCAGACCCCAACTGGTGGAAAGGAGCTTGCCACGGGCAGACCGGCATGTTTCCCCGCAATTATGTC | CACTACAAGTACCGTCGACACGCTCGGGAGGTGGAGCTAGC | ACCCCCGTGAACTAAAAGCTTATTAGCCTGAAGAGCGTCACAGG |
| FPB1-F11 | GAAGCAAAACAAGCATTTAAAGAATTGTTGAAAGAAAAAAGAGTTCCATCTAATGCATCTTGGGAACAAGCAATGAAAATGATTATTAATGATCCAAGATATTCTGCATTGGCAAAATTGTCTGAAAAAAAACAAGCATTTAATGCATAT | ACGGGGCTGCTCTAGAATGGCTAGC | AAGCTTGGCGGTGGCGGGTCTGGTG |
| CI2A-PIN1 | AAAACTGAATGGCCAGAATTGGTTGGTAAATCTGTTGAAGAAGCAAAAAAAGTTATTTTGCAAGATAAACCAGAAGCACAAATTATTGTTTTGCCAGTTGGTACTATTGTTACTATGGAATATAGAATTGATAGAGTTAGATTGTTTGTTGATAAATTGGATAATATTGCACAAGTTCCAAGAGTTGGT | ACGGGGCTGCTCTAGAATGGCTAGC | AAGCTTGGCGGTGGCGGGTCTGGTG |
| BL17-NTL9 | ATGAAAGTTATTTTTTTGAAAGATGTTAAAGGTAAAGGTAAAAAAGGTGAAATTAAAAATGTTGCAGATGGTTATGCAAATAATTTTTTGTTTAAACAAGGTTTGGCAATTGAAGCAACTCCAGCAAATTTGAAAGCATTG | ACGGGGCTGCTCTAGAATGGCTAGC | AAGCTTGGCGGTGGCGGGTCTGGTG |
| VIL1-HP | CATTTGTCTGATGAAGATTTTAAAGCAGTTTTTGGTATGACTAGATCTGCATTTGCAAATTTGCCATTGTGGAAACAACAAAATTTGAAAAAAGAAAAAGGTTTGTTT | ACGGGGCTGCTCTAGAATGGCTAGC | AAGCTTGGCGGTGGCGGGTCTGGTG |
| CSPA-CSD | ATGACTGGTATTGTTAAATGGTTTAATGCAGATAAAGGTTTTGGTTTTATTACTCCAGATGATGGTTCTAAAGATGTTTTTGTTCATTTTTCTGCAATTCAAAATGATGGTTATAAATCTTTGGATGAAGGTCAAAAAGTTTCTTTTACTATTGAATCTGGTGCAAAAGGTCCAGCAGCAGGTAATGTTACTTCT | CGGGGCTGCTCTAGAATGGCTAGC | AAGCTTGGCGGTGGCGGGTCTG |
| CSPB-CSD | GAAGGTAAAGTTAAATGGTTTAATTCTGAAAAAGGTTTTGGTTTTATTGAAGTTGAAGGTCAAGATGATGTTTTTGTTCATTTTTCTGCAATTCAAGGTGAAGGTTTTAAAACTTTGGAAGAAGGTCAAGCAGTTTCTTTTGAAATTGTTGAAGGTAATAGAGGTCCACAAGCAGCAAATGTTACTAAA | ACGGGGCTGCTCTAGAATGGCTAGC | AAGCTTGGCGGTGGCGGGTCTGGTG |
| CKS1 | ATTTATTATTCTGATAAATATGATGATGAAGAATTTGAATATAGACATGTTATGTTGCCAAAAGATATTGCAAAATTGGTTCCAAAAACTCATTTGATGTCTGAATCTGAATGGAGAAATTTGGGTGTTCAACAATCTCAAGGTTGGGTTCATTATATGATTCATGAACCAGAACCACATATTTTGTTGTTTAGAAGACCA | GCTGCTCTAGAATGGCTAGC | AAGCTTGGCGGTGGCGGGTC |

Extended Data Table 3. Sequences and description of the oligos used in this study.

| Oligo pair | Purpose | Foward oligo | Reverse oligo |
|---|---|---|---|
| 1 | amplify GRB2-SH3 library from Twist pool | GCTCGGGAGGTGGAGCTA | TAATAAGCTTTTAGTTCACGGGGGT |
| 2 | amplify PSD95-PDZ3 library from Twist pool | CGGCTCGGGAGGTGGAGCTA | CATTTCGGAGGTCCACACCGTT |
| 3 | amplify 7 domain library from Twist pool | CCGCCACCGCCAAGC | GCTGCTCTAGAATGGCTAGC |
| 4 | linearise the pGJJ046 for GRB2-SH3 library | CTAGCTCCACCTCCCGAG | CGTGAACTAAAAGCTTATTAGTTATGTCACG |
| 5 | linearise the pGJJ068 for PSD95-PDZ3 library | CTAGCTCCACCTCCCGAG | CGGTGTGGACCTCCGAAATG |
| 6 | qPCR plasmid quantification | GCCTACATACCTCGCTCTGC | CAACCCGGTAAGACACGACT |
| 7 | fs primers for PCR1 for GRB2-SH3 aPCA and bPCA libraries | ACACTCTTTCCCTACACGACGCTCTTCCGATCTGGGAGGTGGAGCTAG<br>ACACTCTTTCCCTACACGACGCTCTTCCGATCTHGGGAGGTGGAGCTAG<br>ACACTCTTTCCCTACACGACGCTCTTCCGATCTHHGGGAGGTGGAGCTAG<br>ACACTCTTTCCCTACACGACGCTCTTCCGATCTHHHGGGAGGTGGAGCTAG<br>ACACTCTTTCCCTACACGACGCTCTTCCGATCTNHHYGGGAGGTGGAGCTAG<br>ACACTCTTTCCCTACACGACGCTCTTCCGATCTNHHYYGGGAGGTGGAGCTAG | GTGACTGGAGTTCAGACGTGTGCTCTTCCGATCTGCGTGACATAACTAATAAGC<br>GTGACTGGAGTTCAGACGTGTGCTCTTCCGATCTNGCGTGACATAACTAATAAGC<br>GTGACTGGAGTTCAGACGTGTGCTCTTCCGATCTNNGCGTGACATAACTAATAAGC<br>GTGACTGGAGTTCAGACGTGTGCTCTTCCGATCTHNHGCGTGACATAACTAATAAGC<br>GTGACTGGAGTTCAGACGTGTGCTCTTCCGATCTHWWHGCGTGACATAACTAATAAGC<br>GTGACTGGAGTTCAGACGTGTGCTCTTCCGATCTHWWAAGCGTGACATAACTAATAAGC |
| 8 | fs PCR1 for PSD95-PDZ3 aPCA and bPCA libraries | ACACTCTTTCCCTACACGACGCTCTTCCGATCTGGGAGGTGGAGCTAG<br>ACACTCTTTCCCTACACGACGCTCTTCCGATCTHGGGAGGTGGAGCTAG<br>ACACTCTTTCCCTACACGACGCTCTTCCGATCTHHGGGAGGTGGAGCTAG<br>ACACTCTTTCCCTACACGACGCTCTTCCGATCTHHHGGGAGGTGGAGCTAG<br>ACACTCTTTCCCTACACGACGCTCTTCCGATCTNHHYGGGAGGTGGAGCTAG<br>ACACTCTTTCCCTACACGACGCTCTTCCGATCTNHHYYGGGAGGTGGAGCTAG | GTGACTGGAGTTCAGACGTGTGCTCTTCCGATCTTGACCCGCATTCTTCAG<br>GTGACTGGAGTTCAGACGTGTGCTCTTCCGATCTNTGACCCGCATTCTTCAG<br>GTGACTGGAGTTCAGACGTGTGCTCTTCCGATCTVYTGACCCGCATTCTTCAG<br>GTGACTGGAGTTCAGACGTGTGCTCTTCCGATCTVYMTGACCCGCATTCTTCAG<br>GTGACTGGAGTTCAGACGTGTGCTCTTCCGATCTVYMDTGACCCGCATTCTTCAG<br>GTGACTGGAGTTCAGACGTGTGCTCTTCCGATCTVYMADTGACCCGCATTCTTCAG |
| 9 | fs PCR1 for 7 domains aPCA library | ACACTCTTTCCCTACACGACGCTCTTCCGATCTGCTGCTCTAGAATGGCTAGC<br>ACACTCTTTCCCTACACGACGCTCTTCCGATCTNGCTGCTCTAGAATGGCTAGC<br>ACACTCTTTCCCTACACGACGCTCTTCCGATCTNNGCTGCTCTAGAATGGCTAGC<br>ACACTCTTTCCCTACACGACGCTCTTCCGATCTHWAGCTGCTCTAGAATGGCTAGC<br>ACACTCTTTCCCTACACGACGCTCTTCCGATCTNHTAGCTGCTCTAGAATGGCTAGC<br>ACACTCTTTCCCTACACGACGCTCTTCCGATCTSSSAAGCTGCTCTAGAATGGCTAGC | GTGACTGGAGTTCAGACGTGTGCTCTTCCGATCTCCCGCCACCGCCAAG<br>GTGACTGGAGTTCAGACGTGTGCTCTTCCGATCTNCCCGCCACCGCCAAG<br>GTGACTGGAGTTCAGACGTGTGCTCTTCCGATCTTNCCCGCCACCGCCAAG<br>GTGACTGGAGTTCAGACGTGTGCTCTTCCGATCTNTGCCCGCCACCGCCAAG<br>GTGACTGGAGTTCAGACGTGTGCTCTTCCGATCTGWWWCCCGCCACCGCCAAG<br>GTGACTGGAGTTCAGACGTGTGCTCTTCCGATCTACTWWCCCGCCACCGCCAAG<br>GTGACTGGAGTTCAGACGTGTGCTCTTCCGATCTCATWWWCCCGCCACCGCCAAG<br>GTGACTGGAGTTCAGACGTGTGCTCTTCCGATCTGTATADDCCCGCCACCGCCAAG<br>GTGACTGGAGTTCAGACGTGTGCTCTTCCGATCTTGGGDWWWCCCGCCACCGCCAAG |

Extended Data Table 4. Sequences of indexes used for single- and dual-indexing for Illumina Sequencing.

| Library | Sample | Biological replicate | Technical replicate | FW index primer | FW index | RV index primer | RV index |
|---|---|---|---|---|---|---|---|
| GRB2-SH3 aPCA | input | 1 | 1 | AATGATACGGCGACCACCGAGATCTACACAGAGCCTAACACTCTTTCCCTACACGACGCTCTTC | AGAGCCTA | CAAGCAGAAGACGGCATACGAGATACCAATTAGTGACTGGAGTTCAGACGTGTGCTCTTC | TAATTGGT |
| GRB2-SH3 aPCA | input | 2 | 1 | AATGATACGGCGACCACCGAGATCTACACAGCTATCAACACTCTTTCCCTACACGACGCTCTTC | AGCTATCA | CAAGCAGAAGACGGCATACGAGATACCGAATGGTGACTGGAGTTCAGACGTGTGCTCTTC | CATTCGGT |
| GRB2-SH3 aPCA | input | 3 | 1 | AATGATACGGCGACCACCGAGATCTACACAGGCTCTAACACTCTTTCCCTACACGACGCTCTTC | AGGCTCTA | CAAGCAGAAGACGGCATACGAGATACCTAAGCGTGACTGGAGTTCAGACGTGTGCTCTTC | GCTTAGGT |
| GRB2-SH3 aPCA | output | 1 | 1 | AATGATACGGCGACCACCGAGATCTACACAGGTCGAAACACTCTTTCCCTACACGACGCTCTTC | AGGTCGAA | CAAGCAGAAGACGGCATACGAGATACTGGAGCGTGACTGGAGTTCAGACGTGTGCTCTTC | GCTCCAGT |
| GRB2-SH3 aPCA | output | 2 | 1 | AATGATACGGCGACCACCGAGATCTACACAGTCTGGAACACTCTTTCCCTACACGACGCTCTTC | AGTCTGGA | CAAGCAGAAGACGGCATACGAGATAGAACCGGGTGACTGGAGTTCAGACGTGTGCTCTTC | CCGGTTCT |
| GRB2-SH3 aPCA | output | 3 | 1 | AATGATACGGCGACCACCGAGATCTACACATATTACGACACTCTTTCCCTACACGACGCTCTTC | ATATTACG | CAAGCAGAAGACGGCATACGAGATAGATGCGAGTGACTGGAGTTCAGACGTGTGCTCTTC | TCGCATCT |
| GRB2-SH3 bPCA | input | 1 | 1 | AATGATACGGCGACCACCGAGATCTACACTCTTTCCCTACACGACGCTCTTC | - | CAAGCAGAAGACGGCATACGAGATAAGAACCAGTGACTGGAGTTCAGACGTGTGCTCTTC | TGGTTCTT |
| GRB2-SH3 bPCA | input | 2 | 1 | AATGATACGGCGACCACCGAGATCTACACTCTTTCCCTACACGACGCTCTTC | - | CAAGCAGAAGACGGCATACGAGATCGAATCCAGTGACTGGAGTTCAGACGTGTGCTCTTC | TGGATTCG |
| GRB2-SH3 bPCA | input | 3 | 1 | AATGATACGGCGACCACCGAGATCTACACTCTTTCCCTACACGACGCTCTTC | - | CAAGCAGAAGACGGCATACGAGATCTAGTTGCGTGACTGGAGTTCAGACGTGTGCTCTTC | GCAACTAG |
| GRB2-SH3 bPCA | input | 3 | 2 | AATGATACGGCGACCACCGAGATCTACACTCTTTCCCTACACGACGCTCTTC | - | CAAGCAGAAGACGGCATACGAGATCTAGTTGCGTGACTGGAGTTCAGACGTGTGCTCTTC | GCAACTAG |
| GRB2-SH3 bPCA | output | 1 | 1 | AATGATACGGCGACCACCGAGATCTACACTCTTTCCCTACACGACGCTCTTC | - | CAAGCAGAAGACGGCATACGAGATGATCTTCCGTGACTGGAGTTCAGACGTGTGCTCTTC | GGAAGATC |
| GRB2-SH3 bPCA | output | 1 | 2 | AATGATACGGCGACCACCGAGATCTACACTCTTTCCCTACACGACGCTCTTC | - | CAAGCAGAAGACGGCATACGAGATGATCTTCCGTGACTGGAGTTCAGACGTGTGCTCTTC | GGAAGATC |
| GRB2-SH3 bPCA | output | 2 | 1 | AATGATACGGCGACCACCGAGATCTACACTCTTTCCCTACACGACGCTCTTC | - | CAAGCAGAAGACGGCATACGAGATGCGGTTCTGTGACTGGAGTTCAGACGTGTGCTCTTC | AGAACCGC |
| GRB2-SH3 bPCA | output | 2 | 2 | AATGATACGGCGACCACCGAGATCTACACTCTTTCCCTACACGACGCTCTTC | - | CAAGCAGAAGACGGCATACGAGATGCGGTTCTGTGACTGGAGTTCAGACGTGTGCTCTTC | AGAACCGC |
| GRB2-SH3 bPCA | output | 3 | 1 | AATGATACGGCGACCACCGAGATCTACACTCTTTCCCTACACGACGCTCTTC | - | CAAGCAGAAGACGGCATACGAGATCGACGGTCGTGACTGGAGTTCAGACGTGTGCTCTTC | GACCGTCG |
| PSD95-PDZ3 aPCA | input | 1 | 1 | AATGATACGGCGACCACCGAGATCTACACAACGGCGCACACTCTTTCCCTACACGACGCTCTTC | AACGGCGC | CAAGCAGAAGACGGCATACGAGATAACGACTAGTGACTGGAGTTCAGACGTGTGCTCTTC | TAGTCGTT |

| | | | | | | | |
|---|---|---|---|---|---|---|---|
| PSD95-PDZ3 aPCA | input | 2 | 1 | AATGATACGGCGACCACCGAGATCTACACACCGCGTTACACTCTTTCCCTACACGACGCTCTTC | ACCGCGTT | CAAGCAGAAGACGGCATACGAGATAAGAACCAGTGACTGGAGTTCAGACGTGTGCTCTTC | TGGTTCTT |
| PSD95-PDZ3 aPCA | input | 3 | 1 | AATGATACGGCGACCACCGAGATCTACACACGCTGCAACACTCTTTCCCTACACGACGCTCTTC | ACGCTGCA | CAAGCAGAAGACGGCATACGAGATAAGAAGACGTGACTGGAGTTCAGACGTGTGCTCTTC | GTCTTCTT |
| PSD95-PDZ3 aPCA | output | 1 | 1 | AATGATACGGCGACCACCGAGATCTACACAACGATAGACACTCTTTCCCTACACGACGCTCTTC | AACGATAG | CAAGCAGAAGACGGCATACGAGATAACCGCCAGTGACTGGAGTTCAGACGTGTGCTCTTC | TGGCGGTT |
| PSD95-PDZ3 aPCA | output | 2 | 1 | AATGATACGGCGACCACCGAGATCTACACAACGCCATACACTCTTTCCCTACACGACGCTCTTC | AACGCCAT | CAAGCAGAAGACGGCATACGAGATAACCTCAGGTGACTGGAGTTCAGACGTGTGCTCTTC | CTGAGGTT |
| PSD95-PDZ3 aPCA | output | 3 | 1 | AATGATACGGCGACCACCGAGATCTACACAACGCGCAACACTCTTTCCCTACACGACGCTCTTC | AACGCGCA | CAAGCAGAAGACGGCATACGAGATAACGAAGTGTGACTGGAGTTCAGACGTGTGCTCTTC | ACTTCGTT |
| PSD95-PDZ3 bPCA | input | 1 | 1 | AATGATACGGCGACCACCGAGATCTACACAGAGCCTAACACTCTTTCCCTACACGACGCTCTTC | AGAGCCTA | CAAGCAGAAGACGGCATACGAGATACCAATTAGTGACTGGAGTTCAGACGTGTGCTCTTC | TAATTGGT |
| PSD95-PDZ3 bPCA | input | 2 | 1 | AATGATACGGCGACCACCGAGATCTACACAGCTATCAACACTCTTTCCCTACACGACGCTCTTC | AGCTATCA | CAAGCAGAAGACGGCATACGAGATACCGAATGGTGACTGGAGTTCAGACGTGTGCTCTTC | CATTCGGT |
| PSD95-PDZ3 bPCA | input | 3 | 1 | AATGATACGGCGACCACCGAGATCTACACAGGCTCTAACACTCTTTCCCTACACGACGCTCTTC | AGGCTCTA | CAAGCAGAAGACGGCATACGAGATACCTAAGCGTGACTGGAGTTCAGACGTGTGCTCTTC | GCTTAGGT |
| PSD95-PDZ3 bPCA | output | 1 | 1 | AATGATACGGCGACCACCGAGATCTACACAGGTCGAAACACTCTTTCCCTACACGACGCTCTTC | AGGTCGAA | CAAGCAGAAGACGGCATACGAGATACTGGAGCGTGACTGGAGTTCAGACGTGTGCTCTTC | GCTCCAGT |
| PSD95-PDZ3 bPCA | output | 2 | 1 | AATGATACGGCGACCACCGAGATCTACACAGTCTGGAACACTCTTTCCCTACACGACGCTCTTC | AGTCTGGA | CAAGCAGAAGACGGCATACGAGATAGAACCGGGTGACTGGAGTTCAGACGTGTGCTCTTC | CCGGTTCT |
| PSD95-PDZ3 bPCA | output | 3 | 1 | AATGATACGGCGACCACCGAGATCTACACATATTACGACACTCTTTCCCTACACGACGCTCTTC | ATATTACG | CAAGCAGAAGACGGCATACGAGATAGATGCGAGTGACTGGAGTTCAGACGTGTGCTCTTC | TCGCATCT |
| 7 doms aPCA | input | 1 | 1 | AATGATACGGCGACCACCGAGATCTACACAACGCGCAACACTCTTTCCCTACACGACGCTCTTC | AACGCGCA | CAAGCAGAAGACGGCATACGAGATAACGAAGTGTGACTGGAGTTCAGACGTGTGCTCTTC | ACTTCGTT |
| 7 doms aPCA | input | 2 | 1 | AATGATACGGCGACCACCGAGATCTACACAACGGCGCACACTCTTTCCCTACACGACGCTCTTC | AACGGCGC | CAAGCAGAAGACGGCATACGAGATAACGACTAGTGACTGGAGTTCAGACGTGTGCTCTTC | TAGTCGTT |
| 7 doms aPCA | input | 3 | 1 | AATGATACGGCGACCACCGAGATCTACACACCGCGTTACACTCTTTCCCTACACGACGCTCTTC | ACCGCGTT | CAAGCAGAAGACGGCATACGAGATAAGAACCAGTGACTGGAGTTCAGACGTGTGCTCTTC | TGGTTCTT |
| 7 doms aPCA | output | 1 | 1 | AATGATACGGCGACCACCGAGATCTACACAGAAGCATACACTCTTTCCCTACACGACGCTCTTC | AGAAGCAT | CAAGCAGAAGACGGCATACGAGATACCAAGATGTGACTGGAGTTCAGACGTGTGCTCTTC | ATCTTGGT |
| 7 doms aPCA | output | 2 | 1 | AATGATACGGCGACCACCGAGATCTACACAGAGCCTAACACTCTTTCCCTACACGACGCTCTTC | AGAGCCTA | CAAGCAGAAGACGGCATACGAGATACCAATTAGTGACTGGAGTTCAGACGTGTGCTCTTC | TAATTGGT |
| 7 doms aPCA | output | 3 | 1 | AATGATACGGCGACCACCGAGATCTACACAGCTATCAACACTCTTTCCCTACACGACGCTCTTC | AGCTATCA | CAAGCAGAAGACGGCATACGAGATACCGAATGGTGACTGGAGTTCAGACGTGTGCTCTTC | CATTCGGT |
